# Antibody-mediated immobilization of virions in mucus

**DOI:** 10.1101/500538

**Authors:** Melanie A. Jensen, Ying-Ying Wang, Samuel K. Lai, M. Gregory Forest, Scott A. McKinley

**Author notes:** Submitted to the editors December 15th, 2018. **Funding:** NIH R01GM122082-01, R21AI093242, U19AI096398, NSF DMS-1462992, DMS-1517274, CAREER Award DMR-1151477. **AMS subject classifications**. 92B05, 62-07, 60J70.

## Abstract

Antibodies have been shown to hinder the movement of Herpes Simplex Virus (HSV) virions in cervicovaginal mucus (CVM), as well as other viruses in other mucus secretions. However, it has not been possible to directly observe the mechanisms underlying this phenomenon, so the nature of virion-antibody-mucin interactions remain poorly understood. In this work, we analyzed thousands of virion traces from single particle tracking experiments to explicate how antibodies must cooperate to immobilize virions for relatively long time periods. First, using a clustering analysis, we observed a clear separation between two classes of virion behavior: Freely Diffusing and Immobilized. While the proportion of Freely Diffusing virions decreased with antibody concentration, the magnitude of their diffusivity did not, implying an all-or-nothing dichotomy in the pathwise effect of the antibodies. Proceeding under the assumption that all binding events are reversible, we used a novel switch-point detection method to conclude that there are very few, if any, state-switches on the experimental time scale of twenty seconds. To understand this slow state-switching, we analyzed a recently proposed continuous-time Markov chain model for binding kinetics and virion movement. Model analysis implied that virion immobilization requires cooperation by multiple antibodies that are simultaneously bound to the virion and mucin matrix, and that there is an entanglement phenomenon that accelerates antibody-mucin binding when a virion is immobilized. In addition to developing a widely-applicable framework for analyzing multi-state particle behavior, this work substantially enhances our mechanistic understanding of how antibodies can reinforce a mucus barrier against passive invasive species.

## 1. Introduction

There are several mechanisms by which antibodies (Ab) produced by the immune system can interfere with and even prevent viral infection after an invasion. Antibodies have long been known to bind to surface epitopes on invading virions, rendering the pathogen ineffective either by blocking the epitope from binding to receptors on target cells, or signaling to other immune cells/molecules to inactivate the virus or destroy virus-infected cells. Recent experiments have revealed a previously under-appreciated mechanism: physical *hindrance* of virion motion and potentially the complete *immobilization* of virions in mucus secretions that lie on the epithelium [18, 11]. Specifically, the presence of virion-binding, Immunoglobulin G (IgG) antibody, was shown to directly decrease the mobility of the Herpes Simplex Virus (HSV) virions in human cervicovaginal mucus (CVM) [18], as well as Influenza and Ebola virus-like particles in human airway mucus [22]. An example of the effect can be seen in Figure 1, where we display virion trajectories for two populations of HSV virions, originally studied in Wang et. al. [18]. The left and right columns show virion movement in the presence of low and high Ab concentrations, respectively. The degree of activity in the low Ab concentration is notably higher.

**Figure 1.**
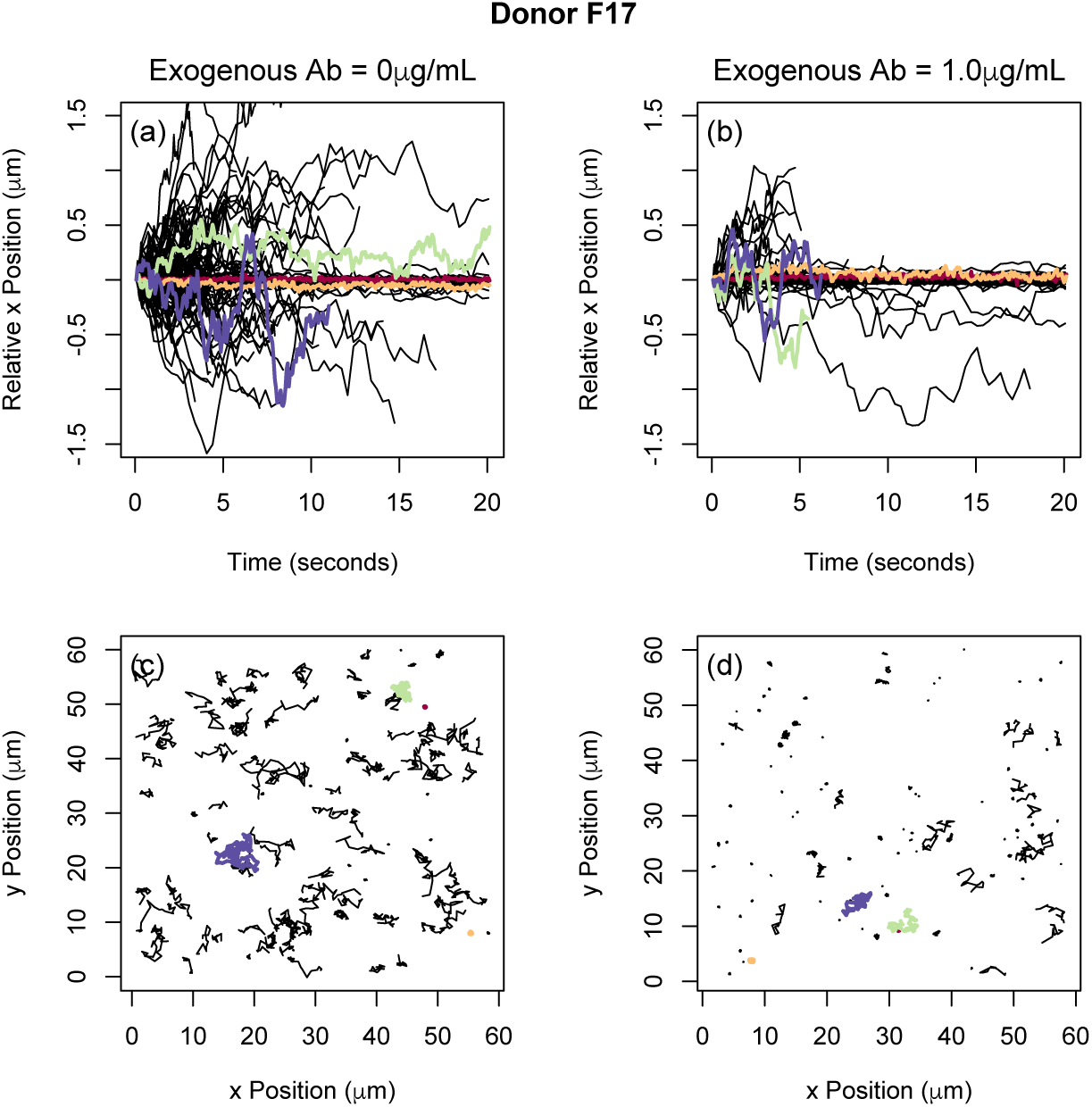
*The trajectories of HSV virions for Donor F17 at exogenous antibody concentrations Oμg/m.L (left) and* 1.0*μg/m.L (right). **Top Row:** The displacement of HSV virions in the x-direction. The time indicated in the horizontal axis is shifted for each path so that t* = 0 *corresponds to the moment the path is first observed. **Bottom Row:** All two dimensional HSV virion trajectories overlaid and plotted in a single frame. For all sub-figures the trajectory frame-rates are 15 observations per second*.

The possibility of using IgG to hinder the motion of different viruses in mucus provides a novel strategy for immunologists to develop methods to prevent and/or treat viral infection [11, 21]. Antibodies are too small to track individually (effective radius ~ 5 nm), but population-scale experimental methods have shown that Ab are slightly less mobile in mucus than in phosphate-buffered saline [13]. The reduced diffusivity of Ab in mucus has been attributed to weak transient bonds between individual Ab and the polymeric microstructure of mucus, or “mucin mesh” [13]. Meanwhile many virions have been shown to diffuse unimpeded in mucus in the absence of a detectable Ab concentration [13, 18]. For this reason, the observation that virion mobility in CVM is impeded in the presence of Ab implies there must be some physico-chemical mechanism at work [18].

Recently the authors and collaborators have explored the possibility that Ab can work in tandem with the mucin mesh to hinder diffusing virions. (See Figure 2 for an idealized schematic of the interactions.) In theory, as a virion diffuses through mucus, an array of Ab can accumulate on its surface. When a sufficient number of virion-bound Ab form low affinity bonds to the mucin mesh, the virion can become tethered and essentially trapped. This hypothesis was introduced by Olmsted et al. in 2001 [13] and confirmed by Wang et al. in 2014 [18] and by Newby et al. in 2017 [11]. In 2014, Chen et al. [2] introduced a stochastic/deterministic hybrid model for the immobilization of Human Immunodeficiency Virus (HIV) by IgG in CVM, and demonstrated the potential impact of the tandem effect of Ab-virion binding and Ab-mucus transient binding on the ability of viral populations to cross, enter, and pass through a thin mucosal layer. Later, Wessler et al. [20] used numerical simulations to explore combinations of Ab-virion and Ab-mucus reaction kinetics that produce an optimal effect. Newby et al. [11] further demonstrated that very low affinity Ab-mucus bonds optimize trapping of diffusing nanoparticles using experimental and simulated data along with providing theoretical arguments.

**Figure 2.**
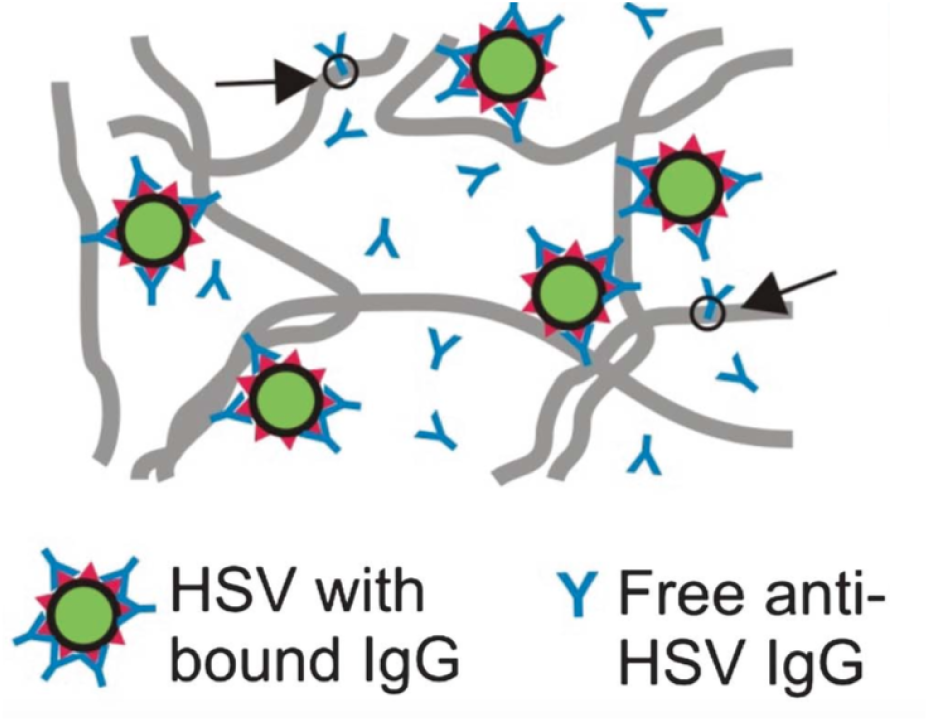
*A schematic depiction of the proposed immobilization process of virions, green circles, by antibodies, blue ‘Y’s, in a mucosal medium. Virions become immobilized, when ‘enough’ antibodies are bound to the virions and the mucosal fibers, gray lines. Arrows indicate Ab interacting solely with the mucin fibers. Figure originally presented in [18]*.

Underlying these mathematical models is a *Switching Diffusion Hypothesis*: that the chemical reactions responsible for virion (or nanoparticle) immobilization are reversible and, as a consequence, virions should switch between diffusive and immobilized states. When compared to the experimentally observable timescale of 10-20 seconds, the Ab-mucin kinetic rates are expected to be fast while the Ab-virion kinetic rates are expected to be slow (see Table 1). It is not clear, however, whether the state-switching between diffusion and immobilization should be on a faster or slower timescale than the observable 10-20 seconds.

**Table 1.** *Parameters and known values incorporated in the model*. * *indicates that the value has not been directly measured. The given value is chosen to be consistent with indirect observations*.

| Parameter | Symbol | Value | Reference |
| --- | --- | --- | --- |
| Cell Properties |  |  |  |
| Initial Ab concentration in CVM | $[A]_0$ | Model Parameter | |
| Concentration of Ab binding sites on mucin fibers in CVM | $[M]$ | unknown | * |
| | $m_{\text{on}}[M]$ | $11.1s^{-1}$ | * |
| Molecule Properties |  |  |  |
| bnAb (IgG) Diameter | | $0.011 \mu m$ | [18] |
| HSV-1 Diameter | | $\sim 0.180 \mu m$ | [18] |
| Number of Ab binding sites on HSV-1 | $N_*$ | Model Parameter | * |
| Reaction Kinetics |  |  |  |
| Ab-mucin affinity (Knockdown Factor) | $\alpha$ | 0.9 | [13] |
| Ab-mucin binding rate | $m_{\text{on}}$ | Unknown | * |
| Ab-mucin unbinding rate | $m_{\text{off}}$ | $100s^{-1}$ | * |
| Ab-virion binding rate | $k_{\text{on}}$ | $4.26e4 [M]^{-1}s^{-1}$ | [2] |
| Ab-virion unbinding rate | $k_{\text{off}}$ | $2.87e-4 s^{-1}$ | [2] |
| Change in (Ab-virion)-mucin binding rate after immobilization | $c$ | Model Parameter | * |
| Number of Ab bond to mucus to immobilize a virion | $T$ | Model Parameter | * |

In multiple papers [2, 20], the number of Ab that were bound to a given virion were tracked and, over the period of time that the number of virion-bound Ab was constant, the virion was assumed to have a state-dependent diffusivity:

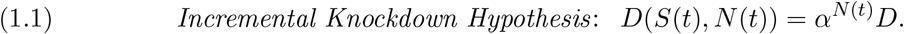

Here *D* is the diffusivity of the virion in mucus in the absence of Ab, *N*(*t*) is the number of virion-bound Ab, and *S*(*t*) is the subset of Ab simultaneously bound to the mucin mesh. This reduction in diffusivity is independent of *S*(*t*) because the number of simultaneously bound Ab changes so rapidly (relative to the number of bound Ab), the virion only feels the average effect of these changes, which is captured by the number of bound Ab, *N*(*t*). The parameter *α* can be expressed in terms of the Ab-mucin binding and unbinding rates (*m*_on_ and *m*_off_, respectively) and the effective concentration [*M*] of binding sites on the surfaces of mucin fibers. If *m*_on_[*M*] and *m*_off_ are very large, so that there are many on-and-off switches per second, then an effective diffusivity arises with a so-called “knockdown factor” *α* = *m*_off_/(*m*_on_[*M*] + *m*_off_) [2]. In this way, we say that the Incremental Knockdown Hypothesis follows from assuming that the dynamics is in a *Fast Switching Regime*. That is to say, in this modeling regime, one assumes that [diffusion ⇄ immobilization] switching is faster than the times between experimental observations and faster than simulation time steps. We depict a typical trajectory of a virion under this hypothesis in Figure 3(a). A virion rapidly changes between the immobilized (red) state and freely diffusing states (green). The resulting path has a reduced *effective diffusivity* that is well-approximated by Equation (1.1), and the virion exhibits qualitatively less movement than a virion predominately in the freely diffusing state (seen in blue).

**Figure 3.**
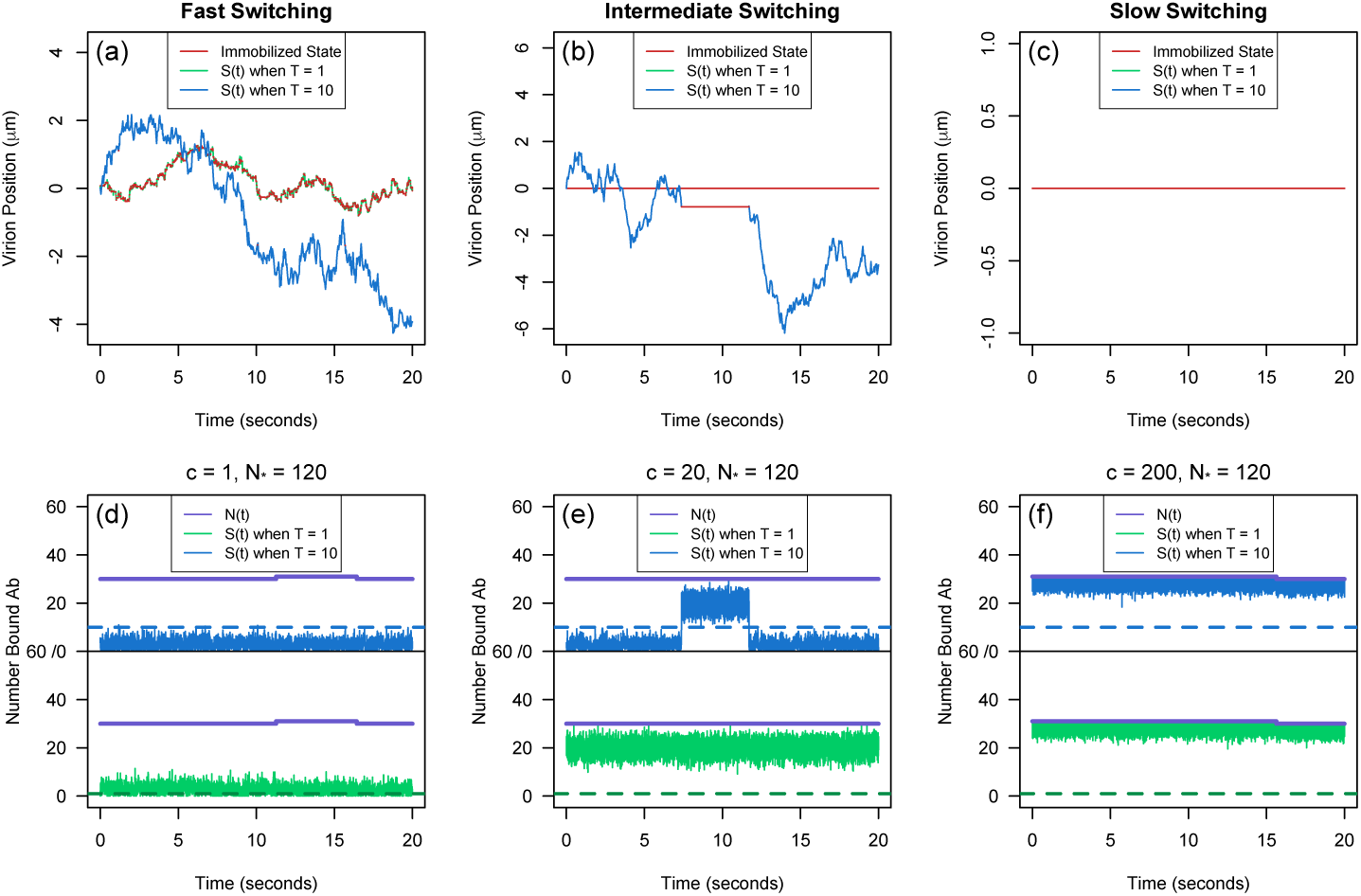
***Top row** (a)-(c): The path of a virion assuming it takes one (green trajectory) or ten (blue trajectory) simultaneously bound Ab for immobilization. Red intervals correspond to periods of immobilization. **Bottom Row** (d)-(f): The virion-Ab-mucin dynamics that govern the movement of the simulated virion directly above it. Within each frame, the number of bound Ab N*(*t*) *is shown by the purple trajectory and the subset of these Ab that are simultaneously bound to the mucin fibers S*(*t*) *assuming a low threshold, T* = 1, *and higher-threshold, T* = 10, *shown by the green trajectory and blue trajectory, respectively. The binding rate cascade factor c increases from, left to right: c* = 1, *c* = 20 *and c* = 200, *respectively. Other m.odel parameters used in the simulation are* ([*A*]_0_, [*A*]_exo_, *N*_*_) = (0.2*μg/mL*, 0.1*μg/mL*, 120). *The mathematical m.odel is fully described in subsection* 2.4.

Recent particle tracking experiments now make it possible to analyze virion behavior as it is modulated by various concentrations of Ab [18]. In Figure 1, we display two populations of HSV virions diffusing in CVM with 0 μg/mL and 1 μg/mL concentrations of exogenous HSV-binding IgG. There is qualitatively less virion movement in CVM with higher concentrations of Ab, but, as we argue below using path-by-path analysis, the trajectories of individual virions appear to resemble either that of a strictly immobilized virion or a strictly freely diffusing virion. This absence of observable switches between immobilized and freely diffusing states might seem to ratify the fast switching hypothesis. However, closer analysis of the freely diffusing particles shows that the diffusivity of freely diffusing virions is essentially the same across all exogenous Ab concentrations. This contradicts the Incremental Knockdown Hypothesis, which predicts the diffusivity should decrease with increasing Ab concentration. While there are essentially no observable switches, and the diffusivity of the free population is not incrementally affected by Ab concentration, we find that the proportion of completely immobilized virions is unmistakably increasing with Ab concentration. (See also [18].) This suggests an alternative hypothesis: we are in a *Slow Switching Regime* where switching takes place fast enough (less than the incubation period of thirty minutes) so that the experiments display different movement patterns, but slow enough (more than twenty seconds) so that we do not see switches in the observational time window.

In this work, we develop and implement the tools necessary for making the preceding claims. To be specific, we use clustering analysis to partition virion paths into a few distinct behavioral patterns. We implement a Bayesian switch-point detection algorithm to assess the prevalence of switches in mobile virions. We develop a Markov chain model for virion-Ab-mucin interactions for use in our characterization of the dependence of virion motility on Ab concentration. A critical feature of this model is the possibility that virion immobilization requires multiple simultaneously surface bound Ab, and that a single virion-Ab-mucin binding event might lead to a cascade of such binding events, which would serve to enhance trapping. Using uncertainty quantification techniques we explore the limitations of the available data, but argue there is a reasonable parameter regime that is fully consistent with experimental observations.

## 2. Data Collection, Statistical Methods, and Mathematical Model

### 2.1. Data collection

Single particle tracking data of HSV virions was collected from seven different CVM samples at five added doses of exogenously anti-HSV-1 IgG, (0, 0.033, 0.1, 0.333, 1.0) μg/mL with an incubation period of half an hour to one hour. For each sample, the virions were tacked for a duration of 20 seconds. The *x*-position and the *y*-position of all traces were observed at a time interval of *δ* = 1/15*s*. For a more detail description of the collection process see the Methods section in [18].

### 2.2. Statistical tools for virion trajectory analysis

We used standard statistical techniques to assess whether the behavior of each virion is consistent with the defining properties of Brownian motion (stationarity with Gaussian independent increments) and to infer physical parameters.

#### 2.2.1. Test for Gaussianity and independence of increments

We used normal quantile-quantile (`qqnorm`) plots to qualitatively verify that the path statistics are approximately Gaussian. The `qqnorm` plots for the increment processes had approximately linear relationships for all particles indicating the *x* and *y* increment processes for all particles could be described as Gaussian.

Noting that if a Gaussian process has uncorrelated increments then the increments are independent, we tested for independence of increments by quantifying the statistical significance of their correlation. Let 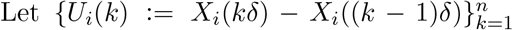 and 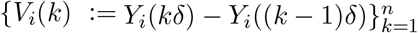 denote the ith particle’s *x* and *y* increment processes, respectively. For the ith particle, we estimated the correlation between the *x* and *y* increment processes separated *h* time steps apart using the sample autocorrelation function, 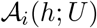 and 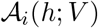 used in the `R` programming language. If there are *n* increments of uniform duration *δ* then for a time lag of *hδ*

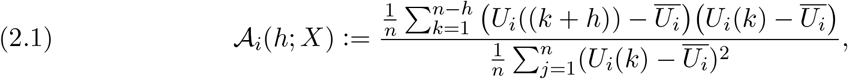

where 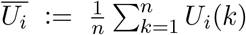 [17]. We say the ith particle’s increment processes are anti-persistent (persistent) if both 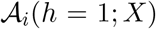 and 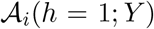 are below (above) the critical value for a 95% significance level and independent otherwise.

#### 2.2.2. Mean-Squared Displacement

The primary statistical tool for describing a population of microparticle paths is the so-called *ensemble* mean-squared displacement (MSD), which we denote 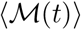. To calculate it, we first compute a *pathwise* MSD for each trajectory (denoted 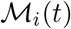 for the *i*th path) and then take an average over these functions. If there are n steps that are uniform of duration *δ*, then as defined in [15],

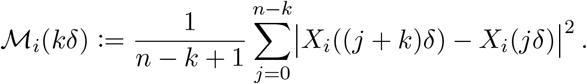

For *t* between the time points {*kδ*} we define 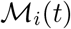 by linear interpolation. The slope of the MSD displayed on a log-log scale provides an estimate for each particle’s diffusive exponent, *ν*, in the large time regime 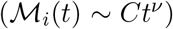. Following standard particle tracking nomenclature, an individual path is said to be Brownian if *ν* = 1, subdiffusive if *ν* ∈ (0, 1), and stationary if *ν* = 0.

#### 2.2.3. Effective Diffusivity

A fundamental quantity to measure for a Brownian path is its *diffusivity D*. If (*X*(*t*), *Y*(*t*)) is the 2*d* position of the particle at time *t*, then its diffusivity is defined to be 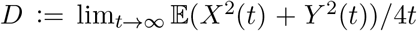. For a Brownian path with *n* steps of uniform duration *δ*, the maximum likelihood estimator (MLE) for its diffusivity has the form

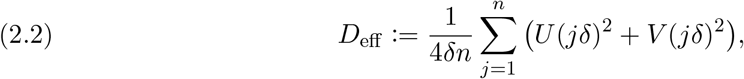

shown in Appendix A. We refer to *D*_eff_ as the path’s *effective diffusivity*. We note that this effective diffusivity is only a consistent estimator for *D* if the path has all the characteristics of Brownian motion, namely stationary, independent, Gaussian increments. However, as seen in Figure 4(a)-(c) there are many paths with anti-correlated increments. For such a process, “diffusivity” is not well-defined. Nevertheless we use *D*_eff_ as a descriptor for these paths because this serves the purpose to distinguish between the particles in two different states by the clustering methods described below.

**Figure 4.**
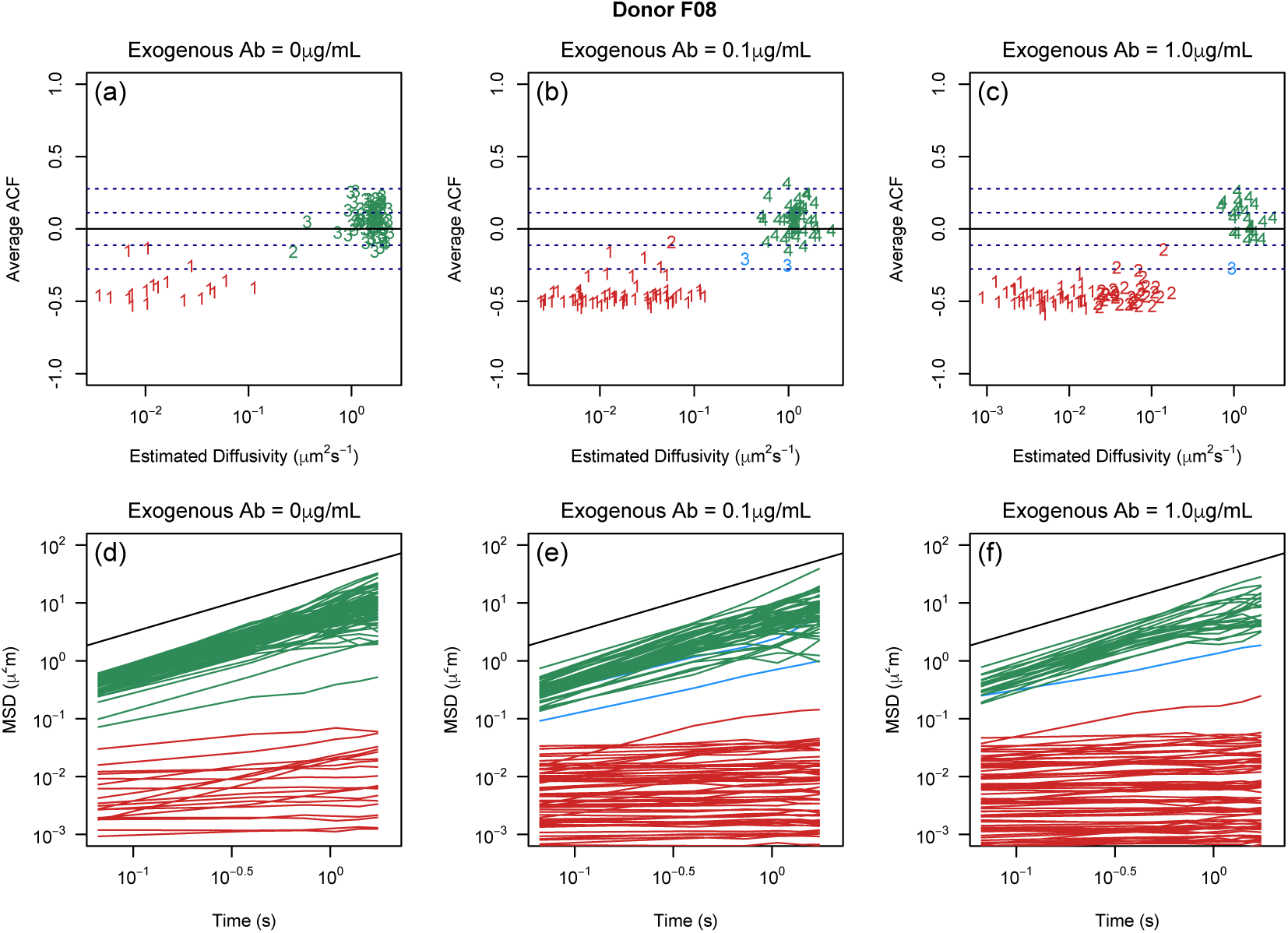
*(a)-(c): The unweighted composition of the tracked virions for Ab concentration 0, 0.1 and 1.0μg/m.L, respectively for Donor F08. Each point corresponds to a tracked virion with the given estimated diffusivity on a log 10 scale and average-ACF value. The character of points denotes clusters prescribed by the hierarchical clustering algorithm, and color of the point denotes the class of the cluster, (d)-(f): The path-wise MSD for all the tracked virions for Donor F08 at [A]_exo_ = 0,0.1, and 1.00μg/m.L. The colors, green, red, and blue, denote the final clusters, freely diffusing, immobilized, and subdiffusive, respectively. Reference line with slope = 1, is denoted in black. (We note that the relative size of the different classes in this figure is not reweighted by path length as it is in the population counts reported in Figure 6.)*

For a given collection of *N* particles, the *ensemble effective diffusivity* is the weighted average effective diffusivity of the tracked particles in the sample, denoted 〈*D*_eff_〉. When evaluating population statistics in particle tracking experiments, if particle paths are weighted independent of path length, then it has been shown that there is a bias toward highly mobile particles, further discussed in subsection 2.3.1, [19]. Based on that analysis, we report the effective diffusivity of an ensemble by taking an average weighted by path lengths. Let 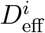 denote the effective diffusivity of ith freely diffusing virion, which has path length *n_i_*. Then

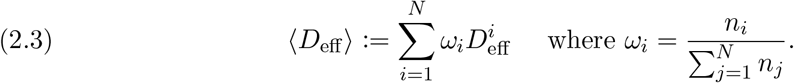

#### 2.2.4 Bias-corrected and accelerated percentile (*BC_a_*) confidence interval method

We constructed confidence intervals for ensemble statistics based on the bootstrapping *BC_a_* method due to its second order accuracy and invariance under transformations. See [4] for the formulation of confidence intervals using this method. We used the boot library in the R programming language to obtain the *BC_a_* confidence intervals for the ensemble statistics as follows. First, we simulated 10,000 booted samples (with replacement) from an ensemble of *N* tracked particles weighted by the particle path lengths. The *BC_a_* confidence interval is then the usual confidence interval constructed using this population of (weighted) bootstrap samples.

### 2.3. Classification scheme for virion paths

For each donor and concentration, we employed a hierarchical clustering algorithm to separate the HSV virions into distinct clusters based on a set of pathwise statistics: *x*-increment ACF, *y*-increment ACF, and log 10 transform of the effective diffusivity. We defined the dissimilarity measure between pairs of virions *i* and *j* by a weighted Euclidean distance *d*(*i*, *j*) with weights of 1/4, 1/4, and 1/2 for the differences in 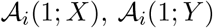, and log10(*D*_eff_) respectively. The dissimilarity measure between clusters was set to be the average linkage. That is to say, the dissimilarity between clusters *R* and *Q* is defined to be

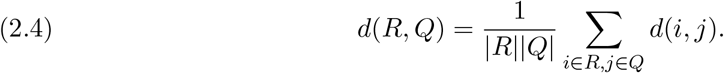

Hierarchical clustering is an agglomerative clustering method [8]. The algorithm is initialized by setting each data point as a distinct cluster. During each iteration, clusters are merged together to minimize the dissimilarity between all clusters. The algorithm stops when all data points are in a single cluster. This process is depicted graphically through the dendrogram where clusters merge at a height equal to dissimilarity between them. We obtained the *k* cluster by cutting the resulting dendrogram at the uniform height yielding *k* clusters.

In almost all cases, we set the number of clusters to *k* = 4 and labeled them Freely Diffusing, Immobilized, Subdiffusive, and Outlier based on cluster ensemble statistics. We introduced an Outlier class to account for those particles whose trajectories were marked by irregular behavior that seemed to be strongly influenced by non-biological factors (likely caused by experimental error). A few examples of each class are displayed in the Supplemental Information, Figure SM1.

The Outlier class and Subdiffusive class were small in number and omitted from the remaining analysis.

#### 2.3.1. ‘Frame-by-frame’ method to compute empirical distribution of each cluster

It has been shown in [19] that fast moving particles are overestimated on shorter time scales in 2d particle tracking. This bias towards the fast moving particles arises due to individual fast particles leaving and reappearing in the focal plane as distinct traces and to new particles entering and leaving the focal plane throughout the duration of the experiment. To minimize overestimating the freely diffusing population, we employed the ‘frame-by-frame’ method developed in [19] to compute the fraction of each population present in the data. The ‘frame-by-frame’ method assigns each tracked particle a weight based on the number of frames the particle appears in the field of view, whereas in the conventional method each particle has the uniform weight of one. Under this weighting system, for a sample of size N, the weighted sample proportion of the *i*th state is given by

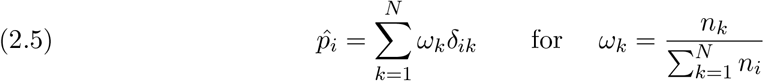

where *δ_ik_* is the Kronecker delta function.

### 2.4. Mathematical model for asymptotic probability of immobilization

We mathematically model the dynamics of a virion under the Switching Diffusion Hypothesis by the following SDE:

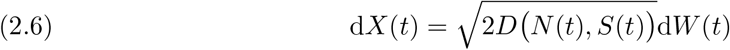

where *W*(*t*) is standard 2d Brownian motion and the state-dependent diffusivity, *D*(*N*(*t*), *S*(*t*)), depends on two time-dependent processes: *N*(*t*), the number of antibodies bound to the surface of a focal virion at time *t*, and *S*(*t*), the subset of these antibodies simultaneously bound to mucin binding sites at time *t*. We establish a threshold parameter T. A virion is defined to be *immobilized* if there are at least *T* simultaneously bound antibodies, *S*(*t*) ≥ *T*, and defined to be *freely diffusing* if there are fewer than *T* simultaneously bound antibodies, *S*(*t*) < *T*. Under this convention, the time-dependent diffusivity is given by

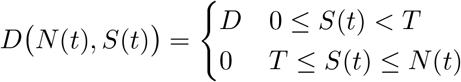

where the constant *D* is the diffusivity of the virion in mucus in the absence of Ab. In the following sections, we present a mathematical model that describes the asymptotic probability of the immobilized state when exposed to varying exogenous antibody concentrations.

### 2.4.1. Model assumptions

Based on the initial population clustering analysis, there appears to a subpopulation of virions that do not interact with the antibodies. We define *q* to be the probability that a given virion will interact with the Ab population. Second, for the sake of simplicity, we assume that Ab-virion binding sites operate independently from each other. However, we allow for conspiracy among the Ab in binding to the mucosal environment. Once the virion has T simultaneously bound Ab-mucin-virion interactions (*S* ≥ *T*) the surface bound antibodies might bind to the mucin fibers differently than if the virion was freely diffusing. We parametrize this by a multiplicative change in Ab-mucin binding rate through the introduction of the dimensionless parameter *c*. If *c* > 1 the parameter has a cascade effect, aiding in the immobilization process [3, 6, 7, 9, 11].

### 2.4.2. A Markov Chain model for virion-Ab-mucin dynamics

Let *N*_*_ denote the number of independent Ab binding sites on the surface of an HSV virion. Antibodies bind and unbind from these sites at rates *k*_on_ and *k*_off_, respectively, with dissociation constant *k*_d_:= *k*_off_/*k*_on_.

Virion-surface-bound antibodies interact with the surrounding mucosal medium, binding to and unbinding from mucin binding sites, at rates *m*_on_ and *m*_off_, with dissociation constant *m*_d_:= *m*_off_/*m*_on_. The total Ab concentration [*A*] is the sum of the exogenous [*A*]_exo_ and endogenous [*A*]_0_ Ab concentrations, and the total concentration of binding sites on mucin fibers is denote [*M*]. See Table 1 for a comprehensive list of variables.

We model the the Ab-virion interactions using a continuous time Markov Chain (CTMC) assuming linear state transitions. If a given virion has *n* occupied (Ab-bound) surface binding sites at time *t*, then the CTMC transition rates are given by

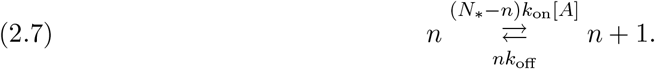

If there are *s* simultaneously bound Ab cross-linking the virion to mucin fibers at time *t* and *n* occupied virion-surface-binding sites, then the conditional Ab-mucin dynamics are modeled by a CTMC with state transition rates

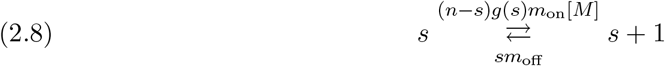

for *s* ≤ *n*, where

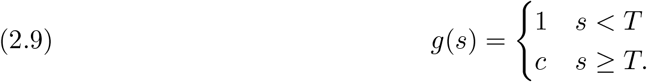

The function in Equation (2.9) quantifies the impact immobilization has on the rate at which additional antibodies crosslink to the mucin fibers, i.e. the binding cascade effect, and results in a non-linear transition rate when *c* ≠ 1. We note that the transition (*n*, *s*) → (*n* − 1, *s* − 1) is omitted from our analysis to facilitate with explicit likelihood calculations. This does not qualitatively affect our results.

We show the impact the immobilization threshold, *T*, and the cascade factor, *c*, have on the immobilization process in Figure 3. Within each frame, it can be seen that a higher immobilization threshold allows for longer freely diffusion periods, while across frames a higher cascade factor leads to longer immobilized periods. In Figure 3(d)-(f) we simulated realizations of the processes (*N*(*t*), *S*(*t*)) for various combinations of *T* and *c*. The number of bound antibodies, *N*(*t*), is displayed by the purple trajectory, and the number of simultaneously bound Ab with a low immobilization threshold, *S*(*t*) when *T* = 1, and with a higher immobilization threshold, *S*(*t*) when *T* = 10, are shown by the green and blue trajectory, respectively. Moving left to right, the factor by which the Ab-mucin binding rate changes after immobilization increases, *c* = 1, 20, and 200, respectively. In Figure 3(a)-(c), we show how these processes dictate the movement of the virion. The virion with process (*N*(*t*), *S*(*t*) when *T* = 1) is colored in green while (*N*(*t*), *S*(*t*) when *T* = 10) is colored in blue. For both trajectories immobilized periods, *S*(*t*) ≥ *T*, are colored in red.

When immobilization does not affect the Ab-mucin binding rate, Figure 3(d), the process S(*t*) rapidly crosses the immobilization threshold (dashed line) resulting in a virion transitioning between states faster than the experimental time step, Figure 3(a), for both *T* = 1 and *T* = 10. By increasing the cascade factor, Figure 3(e)-(f), *S*(*t*) remains above the immobilization threshold, for observable periods. In this case, the simulated virions in Figure 3(b)-(c) change states on the experimental time scale of twenty seconds and longer than twenty seconds, respectively.

### 2.4.3. Our approximation for the stationary probability of being immobilized

We assume that the antibody-virion dynamics are slow compared to the antibody-mucin dynamics. To approximate a virion’s long-term probability of being immobilized, we use a product of two factors. The first is the steady-state distribution for the number of surface-bound Ab, *N*(*t*). Then we compute the stationary distribution for the number of simultaneously bound Ab, *S*(*t*), conditioned on each value *N*(*t*) = *n* (where *n* ∈ {0, … *N*_*_}).

We introduce the notation *b*(*x*, *n*, *p*) for the binomial probability mass function. That is, if *X* ~ Binom(*n*, *p*), then 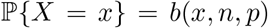. Our approximation to the stationary distribution of immobilization can be understood as an average over the transitions of the fast process *S*(*t*). Let *σ* denote the time a particle spends in the immobilized state, and *τ* the time a particle spends in the freely diffusing state. Then our approximation takes the form

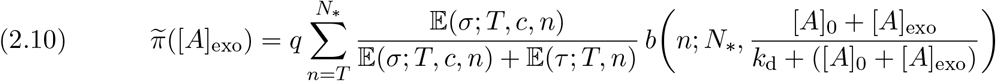

where

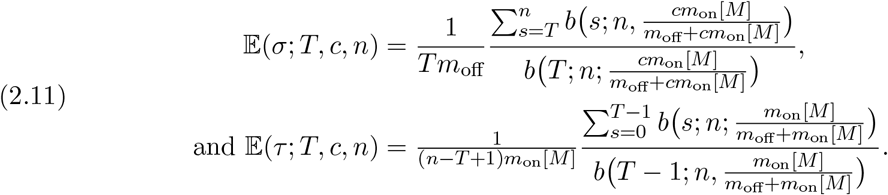

The derivation of Equation (2.10) and Equation (2.11) rely on Markov Chain Theory and Renewal Theory and can be found in Appendix B.

It follows from the law of total expectation and the time-scale approximation, the expected time immobilized and expected time freely diffusing are respectively:

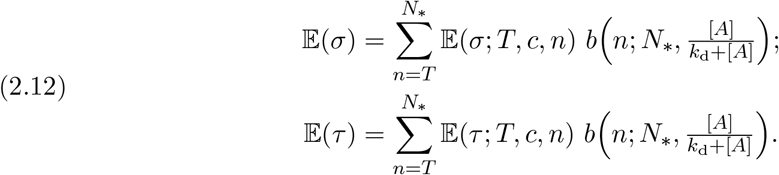

We say that a parameter vector is in the *Slow Switching Regime* if, for all tested exogenous Ab concentrations, the average times spent in the immobilized and diffusing states are more than 20 seconds. To be precise, we define

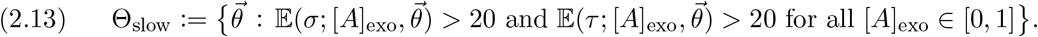

## 2.5. Switch point detection

We develop an algorithm for detecting whether there is a *single* switch from diffusion to immobilization or immobilization to diffusion. The mathematical model presented in subsection 2.4, Equation (2.6), assumes complete immobilization but in fact immobilized virions exhibit spatial motion. Bernstein and Fricks in [1] account for this spatial motion by describing the bound state as a diffusing particle trapped in a potential well. Using an Expectation-Maximization algorithm they provide an evolving probability for each particle that it is in an immobilized or diffusing state. In contrast to the many-switch paths considered by Bernstein and Fricks, we argue in subsection 3.1.2 that the virion paths in our data set have at most one or two switches. We therefore developed and implement a Bayesian algorithm that is designed to identify the presence of a single switch point.

To derive a likelihood function, we extend our SDE model Equation (2.6) to include a path-specific trapping potential well, similar to [1]. Our extended model for a [diffusion → immobilization] switch is

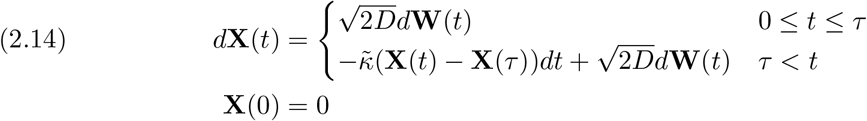

and for [immobilization → diffusion], we have

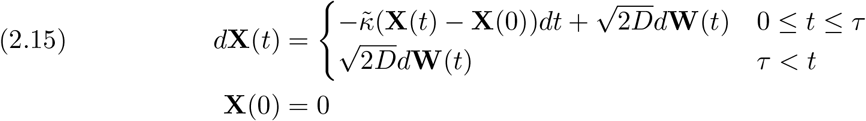

where **X**(*t*) = (*X*(*t*), *Y*(*t*))^*T*^ and **W**(*t*) is 2d Brownian Motion. These SDEs are derived from the Langevin equation for particles diffusing in a quadratic (Hookean spring) potential well. The constant 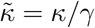 where *κ* is the spring constant and *γ* is the viscous drag experienced by the particle. Due to the Fluctuation-Dissipation relationship, *γ* also appears in the diffusivity constant, which has the form 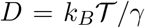, where *k_B_* is Boltzmann’s constant and 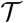 is the temperature of the fluid. To obtain an analytically trackable likelihood function, we introduce simplifying assumptions that (1) the switch occurs exactly at an observation time point, and (2) there is no measurement error. We derive the likelihood function in Appendix C.

We take a Bayesian approach to jointly estimate *D*, 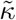, and *τ* under both switching scenarios using a Gibbs sampling algorithm. If the 95% credible region for *τ* is completely contained within the interval [0.1T_final_, 0.9T_final_] where T_final_ is the duration of a path, then we say that path is a candidate for switching. For both switching scenarios we estimated a false discovery rate for this criterion by simulating freely diffusing particles and setting the false discovery rate to the percent of simulated Brownian particles that were labeled as candidates for switching for the given switching model, Equation (2.15) or Equation (2.14). Similarly, we estimated the power of criterion through simulation. For both scenarios we simulated particles that switched states once, and set the power to the fraction of paths that were candidates for switching. See section SM4 for more details on how these tests were constructed, and the results are presented in subsection 3.1.2.

## 2.6. Uncertainty quantification

The model given by Equation (2.10) depends on the parameter vector 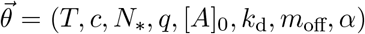. In specifying the model to HSV-IgG data, subsection 2.1, we set *k*_d_ = 0.8969 [10] and *α* = 0.90 [13]. The Ab-mucin binding and unbinding rates have not been directly estimated. We assume they are fast compared to the experimental time scale and, for example, set *m*_off_ = 100*s*^−1^. To assess the remaining parameters, 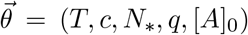 – which are are the immobilization threshold value, the binding cascade factor, the number of sites on the surface of virions, the virion-Ab interaction probability, and the endogenous Ab concentration – we employed the numerical method of profile likelihoods [5, 16]. We used the numerically obtained relationships among parameters to obtain conditions on 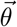 such that the Switching Diffusion Hypothesis (in the Slow Switching Regime) is consistent with the data.

In order to quantify the model’s error in predicting the immobilized fraction, for each donor *i*, we partitioned the paths according to exogenous Ab concentration 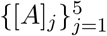, and introduced the following residual function:

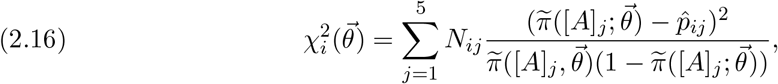

where 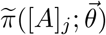 denotes the model evaluated at [*A*]_*j*_ with parameters 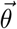 (as defined in Equation (2.10)), while *N_ij_* and 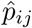 are, respectively, number of paths observed and the fraction that are immobilized in the *j*th subpopulation associated with donor *i*. Assuming a normal approximation to the binomial distribution, our residual function can be seen as the sum of five independent squared normal random variables, i.e. with a *χ*^2^-distribution with 5 degrees of freedom.

### 2.6.1. Numerical method of profile likelihoods to deduce parameter identifiabilty

Because we assume normal approximation to the binomial distribution, working with a residual function is equivalent to using a likelihood function to define confidence intervals [14, 16]. For ease of notation in this section, we will suppress the dependence on *i* when considering the residual function 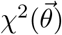 for donor *i*.

To discuss identifiability of our model parameters, we use the nomenclature introduced by Raue in [16]. Our minimum residual estimator is defined to be 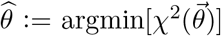. The *likelihood-based confidence region of level α* for 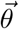 is then defined to be

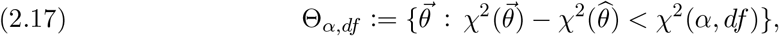

where *χ*^2^(*α*, *df*) is the *α* quantile of the *χ*^2^ distribution with df degrees of freedom. When establishing a confidence interval for one of the parameters, we set *df* = 1. When establish a confidence region for multiple parameters, we set df equal to the number of parameters [14].

A parameter *θ_k_* is said to be *structurally identifiable* when there is a unique minimum of 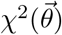 with respect *θ_k_*, i.e., if there exists a unique *θ_k_* such that

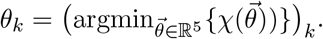

Alternatively, *θ_k_* can be unidentifiable due to the structure of the model or because the quality and quantity of the data is insufficient in estimating *θ*_*k*_. For the former case, we say *θ_*k*_* is *structurally unidentifiable* if the set

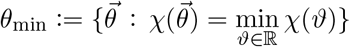

is not unique and contains at least two elements whose *θ_k_* components are distinct. This often occurs when there is a functional relationship *ϕ* among *θ_k_* and at least one other parameter, say *θ_j_* such that % can be expressed directly in terms of *ϕ*(*θ_k_*, *θ_j_*). As for the latter data-restricted type of unidentifiability, we say *θ_k_* is *practically unidentifiable* when a unique minimum exists of 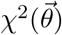 with respect *θ_k_* but the likelihood based confidence interval for 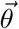 extends infinitely in increasing and/or decreasing values of *θ_k_*.

These definitions can be interpreted graphically using profile likelihoods. For residual function 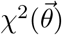 the *profile likelihood* of the *k*-th parameter defined to be

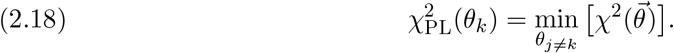

If *θ_k_* is a structurally identifiable parameter then 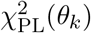 exceeds the threshold Δ*_α_* for both increasing and decreasing values of *θ_k_* forming a deep valley around 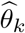. If *θ_k_* is structurally unidentifiable the profile likelihood is flat. Lastly, if *θ_k_* is practically unidentifiable, 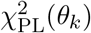 obtains a unique minimum but does not exceed Δ*_α_* in increasing and/or decreasing values of *θ_k_*, forming a shallow valley around 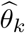.

We further investigate unidentifiable combinations of parameters by extending Equation (2.18) to profile parameter *θ_j_* and *θ_k_* simultaneously,

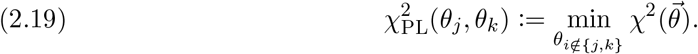

Structural relationships between the two profile parameters manifest as flat valleys extending infinitely along the functional relationship in the contour plots of 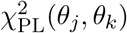. We note this flat valley only traces out the functional relationship *θ_j_* and *θ_k_* when the dimension of the parameter space is larger than 2.

## 3. Results

### 3.1. Data do not support the Incremental Knockdown Hypothesis for a 20 second time frame

#### 3.1.1. No evidence of fast switching: ensemble effective diffusivities of the free subpopulation are the same regardless of exogenous Ab concentration

For each Donor/Ab-concentration combination, the associated sample of virions contained a clear division among the tracked particles’ MSD and ACF behavior. We used the classification scheme described in subsection 2.3 to label each tracked virion as Immobilized, Freely-Diffusing, Subdiffusive or Outlier. The Immobilized class was characterized by low effective diffusivity (< 10^−1^*μm*^2^*s*^−1^) and either anti-persistent or uncorrelated increment processes. Meanwhile the Freely-Diffusing class had uncorrelated increment processes and effective diffusivities larger than 0.2*μm*^2^/s. The Subdiffusive and Outlier classifications were rare and did not appear in all samples. For this reason, we removed these categories from the analysis but give a description of them in the SI. In Figure 4(a)-(c), we display the results of the classification for Donor F08 at 0, 0.1, and 1 *μ*g/mL added anti-HSV IgG in terms of *D*_eff_ and the average of the *x*- and *y*-ACF, as defined in subsection 2.2.3 and subsection 2.2.1 respectively. The clear separation of groups and locations of the clusters were qualitatively similar for the other donors (further figures included in the Supplemental Information).

The pathwise MSDs for Donor F08 virions are displayed in Figure 4(d)-(f), and we note the similarity of the Freely-Diffusing category of virions across all three panels. The Incremental Knockdown Hypothesis would predict that freely diffusing virions would be “slower and slower” in the presence of more and more Ab. However, we found that the diffusivit.ies of the Freely-Diffusing classes are consistent across all exogenous Ab concentrations. In Figure 5 we display this fact in two ways. In the left panel, we display the ensemble MSD averaged over the Freely-Diffusing (green triangles) and Immobilized (red x’s) populations for each Ab concentration. There is remarkable overlap within each group. Moreover, in the right panel, we display the ensemble effective diffusivity for the Freely-Diffusing class at the various exogenous Ab concentrations for all donors. While there is variation in the effective diffusivity, the overlapping *BC_a_* confidence intervals indicate there is insufficient evidence to conclude the effective diffusivity decreases with antibody concentration. (We provide 95% weighted bootstrap confidence intervals for each estimate in the supplementary material Figure SM12).

**Figure 5.**
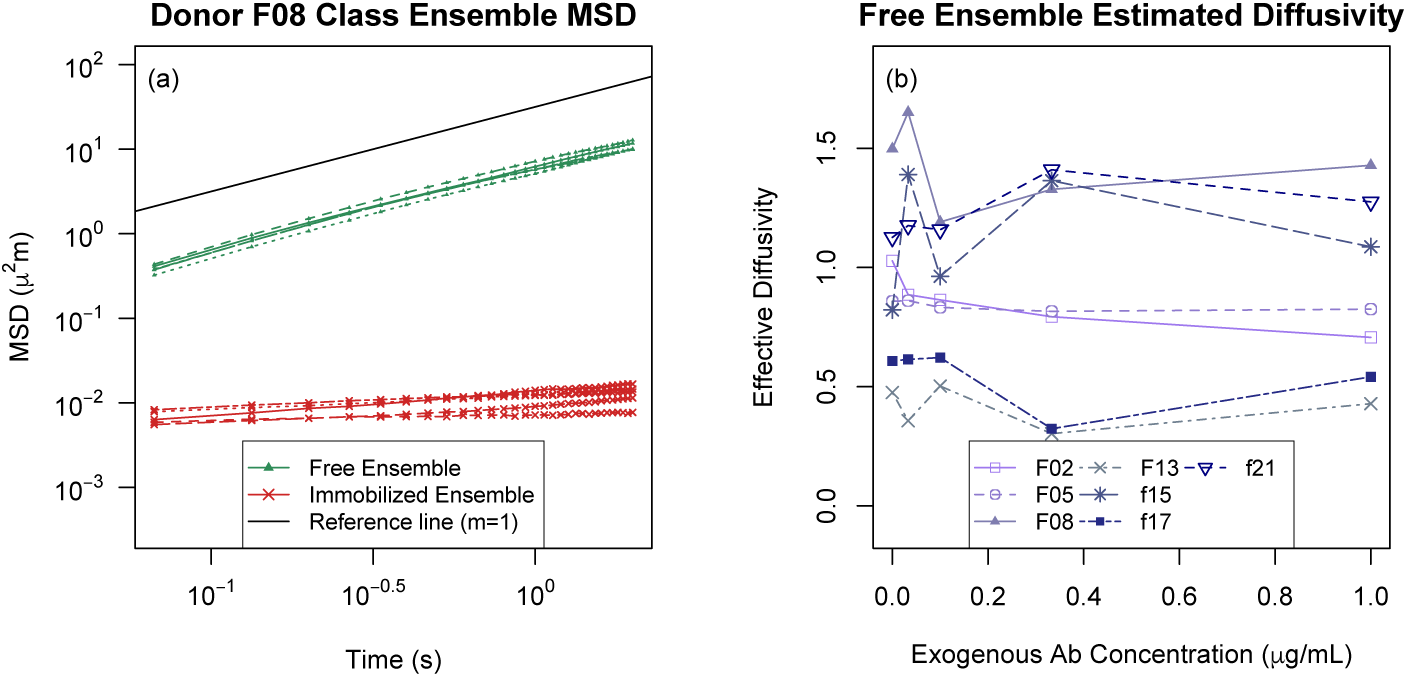
*(a) Ensemble MSD of the Freely Diffusing class and the immobilized class at various exogenous antibody concentrations represented by the green and red curves, respectively, for Donor F08. The black line refers to the ensemble MSD of Brownian particles, slope equal to 1. (b) The estim.ated ensemble effective diffusivity of the free population versus exogenous antibody concentration where the shade and point style of the curve corresponds Donor. See Figure SM12 for the ensemble effective diffusivity with 95% BC_a_ confidence intervals*.

We can express this finding in terms of a statistical test by comparing the weighted ensemble effective diffusivity for the freely diffusing subpopulation at the two extreme Ab concentrations. We used a one-tailed paired difference hypothesis test:

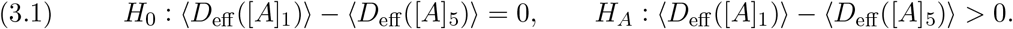

for [*A*]_1_ = 0.0*μ*g/mL and [*A*]_5_ = 1.0*μ*g/mL. At an *α* = 0.05 level of significance, we failed to find significant evidence that the ensemble effective diffusivity of the freely diffusing population decreased when exogenous Ab concentration increased from zero exogenous Ab to the highest concentration (*t*_6_ = 0.2567, *p*-value= 0.4030). We report the results of paired difference tests for all other combinations of the tested exogenous Ab concentration in Table SM10.

#### 3.1.2. Little evidence of switching on the experimental time scale

We found little evidence that virions switch between states on the experimental time scale of 20 seconds. If tracked particles were typically experiencing many subtle switches, we expect that their computed effective diffusivities would be diminished by a factor determined by the time spent immobilized. Moreover, because there are distinct behavioral regimes, the distribution of the increment processes are essentially a mixture of two Gaussian distributions (one for the Immobilized state and one for the Freely Diffusing state). This would manifest itself as a violation of linearity in `qqnorm` plots, which we do not see for the vast majority of HSV virion paths.

While the `qqnorm` test can identify paths that might experience switches, they do not affirm the presence of a switch. To this end, we developed a Bayesian method for identifying whether there is a single switch point in a given virion path, described in the subsection 2.5.

We say a path of duration T_final_ is a candidate for switching if the 95% credible region for *τ* was completely contained within the interval [0.1T_final_, 0.9T_final_]. The method was very effective on simulated data. When we applied the method to simulated Brownian motion (Freely-Diffusing), we found a 0.0119 and 0.0080 False Discovery Rate of [diffusion → immobilization] switches and [immobilization → diffusion] switches, respectively. On the other hand, 96.38% of the simulated [diffusion → immobilization] paths were correctly identified as [diffusion → immobilization] switches, while 94.37% of the simulated [immobilization → diffusion] paths were identified as [immobilization → diffusion] switches (Table 2). Under this method, we found that 1.12% of the Freely-Diffusing class (1689 total tracked virions) were identified as [diffusion → immobilization] switch candidates and 1.24% of the free populations were [immobilization → diffusion] switch candidates. We therefore concluded that state switches occurred relatively rarely on the experimental time scale.

**Table 2.**
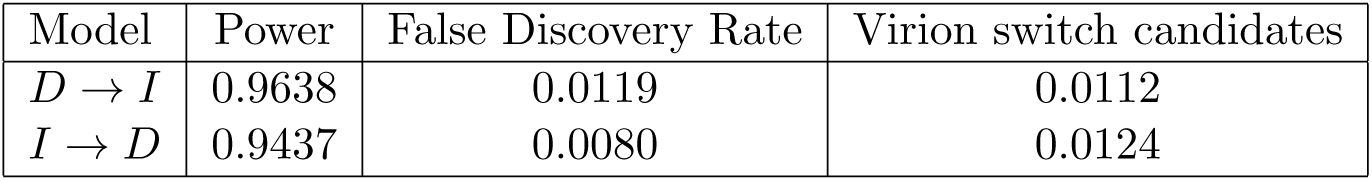
*Fraction of Freely Diffusing virions that possibly switched states once by Donor*.

#### 3.1.3. Fraction immobilized increases with exogenous antibody concentration

While Ab concentration did not seem to affect the behavior of virions labeled Freely-Diffusing, it did have a significant effect on the fraction of virions that were placed in this class. This is consistent with the findings reported in [18]. We computed the Immobilized fraction for each Donor/Ab-concentration sample using the method discussed in subsection 2.3.1 and display the results in Figure 6, where each curve in the panel (b) corresponds to a different donor. While there is heterogeneity in the fraction of Immobilized virions across donors, there is a visible overall increase in proportion immobilized from 0 to 1 *μ*g/mL. This qualitative assessment is supported by statistical evidence provided by non-overlapping *BC_a_* confidence intervals between the extreme exogenous Ab concentrations Figure SM11.

**Figure 6.**
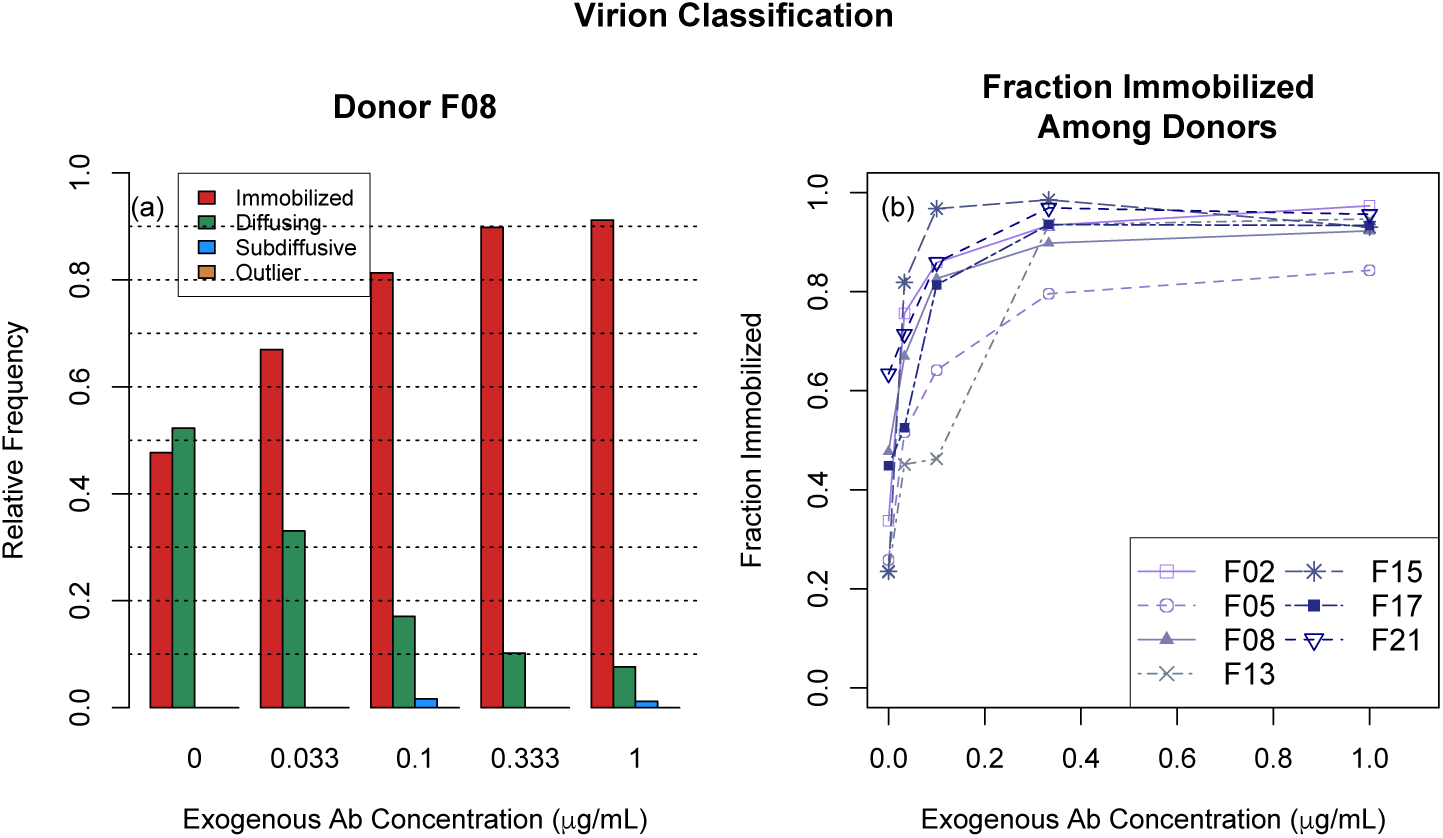
***(a)** The weighted proportion of the 4 classes for Donor F08 at the various tested exogenous Ab concentrations, **(b)** Weighted proportions of Immobilized virions for each donor. See Figure SM11 for plots with 95% BC_a_ confidence intervals*.

For each donor, the fraction of Immobilized virions increased with Ab concentration in the 0 to 0.333 *μ*g/mL range and seemed to be saturated at higher Ab concentrations. We tested the significance of this observed trend by fitting a negative exponential growth model with predictors: exogenous antibody concentration and individual effect terms relative to Donor F08. Let *χ_k_* be the the indicator function that a virion in the *k*th donor sample is in the immobilized state. Our negative exponential growth model takes the form

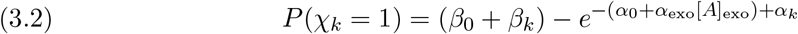

where *α_k_* and *β_k_* are the effect terms for the *k*-th donor. We found the exogenous antibody concentration (*α*_exo_ = 15.920, *p*-value< 0.001), the growth rate due to the baseline donor (*α*_0_ = −0.8427, *p*-value = 0.0043), and baseline saturation probability (*β*_0_ = 0.9138, *p*-value< 0.0001) were statistically significant in predicting the immobilization probability, whereas the constants accounting for deviations from the baseline due to donor sample were not significant. The model was fit using the `R` command *nls*() with the minimization algorithm set to Gauss-Netwon’s method.

### 3.2. The Simple Linear Model predicts fast switching

The results from subsection 3.1 provide evidence against the hypothesis that switching between the diffusing and immobilized states is fast relative to the experimental time scale. Our next goal was to determine whether there is a parameter regime that predicts slow switching while simultaneously being consistent with the exogenous Ab-dependent, Immobilization data displayed in Figure 6. This analysis depends strongly on two assumptions: (1) whether one virion-bound Ab is sufficient to crosslink the virion to mucin, and (2) whether Ab-mucin binding rates increase when the virion is immobilized, the so-called cascade effect. We introduced two variables – *T*, the threshold number, and *c*, the cascade factor – in our general model to account for these possible effects. In recent works, it has been assumed either that *T* = *c* = 1 [2, 11] or that *T* = 1 and *c* > 1 [20]. We refer to *T* = *c* = 1 as the Simple Linear Model (SLM) because all the CTMC transition rates are linear. By computing the expected durations of the immobilized and diffusing states (Equation (2.12), derivation in Appendix B.2), we were able to show that the data is not consistent with the SLM, or any case where *T* = 1.

We say a model is *consistent* with the observed data for a specified donor if there exists a parameter vector 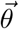 that is within the 95% confidence region for the Immobilized Fraction data (denoted Θ_*α,df*_, defined in Equation (2.17)) and also predicts expected state times larger than 20 seconds (denoted Θ_slow_, defined in Equation (2.13)). In Figure 7, we demonstrate that the SLM is not consistent with the data for Donor F08. In the left panel, we show a 2d profile likelihood plot, for the endogenous Ab concentration [*A*]_0_ and number of virion surface binding sites *A*\*. For each ([*A*]_0_, *N*\*) pair, we calculated the best fit for the remaining parameter *q*, the virion-Ab interaction probability, and display the residual value by the shading (darker means better fits). The black region represents the 95% confidence region for these two parameters.

**Figure 7.**
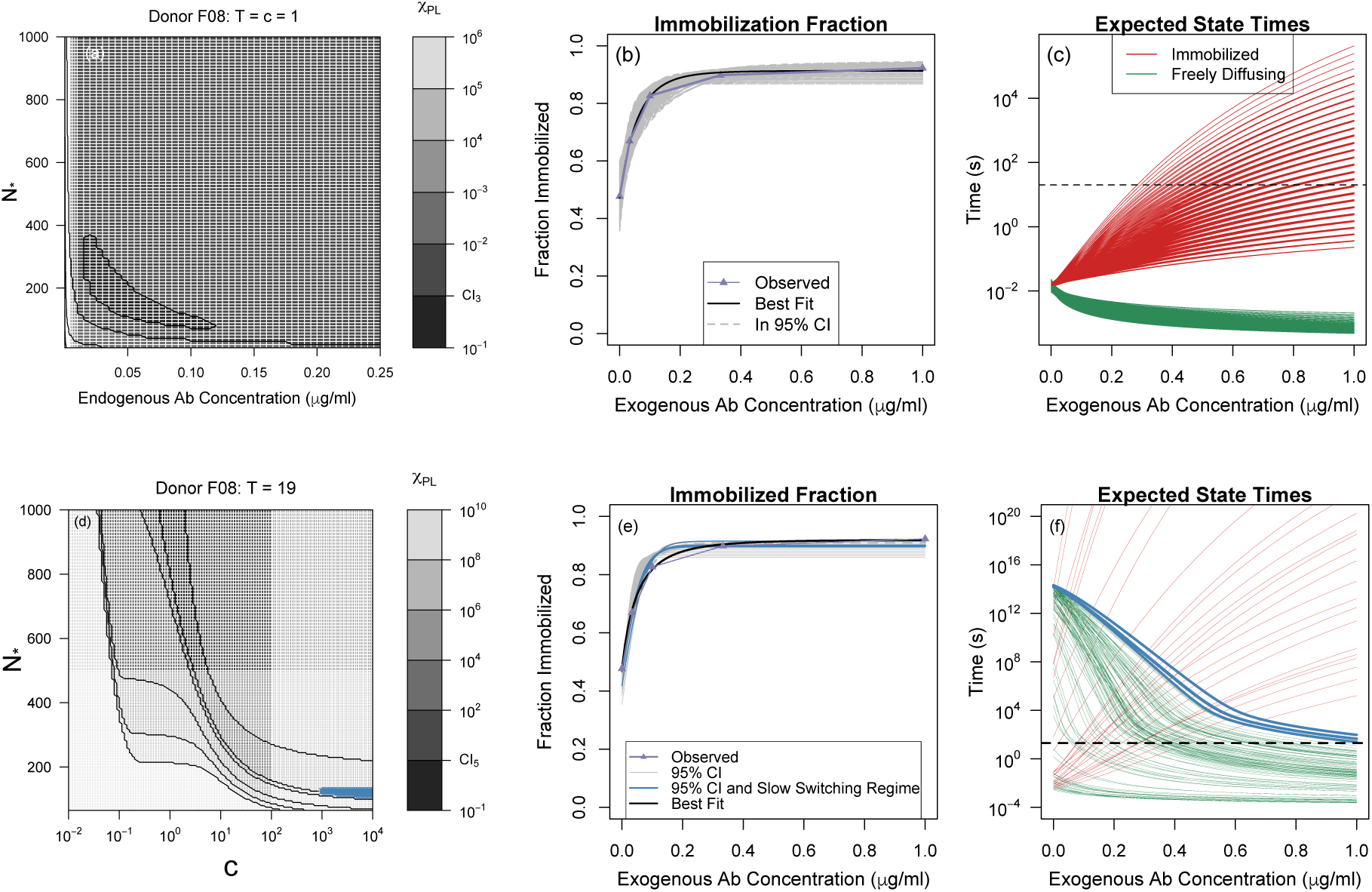
***(a),(d)** Profile likelihood contour plots (Donor F08) for 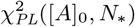 and 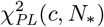 when T = c = 1 and T = 19, respectively. Darker shades correspond to smaller profile likelihood values and the black region corresponds to the 95% confidence regions Θ_0.05,3_ and Θ_0.05,5_. **(b),(e)** Predicted Immobilized Fraction curves (gray lines) for θ sam.pled from, Θ_0.05,3_ and Θ_0.05,5_. The black curve is the prediction of the best fit in each case for Donor F08. The observed Immobilized Fraction is shown by the purple line with. triangles, **(c),(f)** Expected duration of Immobilized (red curves) and Freely-Diffusing (green curves) states for θ sam.pled from. Θ_0.05,3_ and Θ_0.05,5_. When T = c = 1, frame (c), none of predicted state tim.es are above 20 seconds, horizontal black line. On. the other hand, when T = 19, frame (f), there are som.e parameter combinations that do yield slow switching. These are marked in light blue as appropriate in Panels (d)-(f)*.

We uniformly sampled this confidence region, Θ_0.05,3_, and displayed the predicted Immobilized Fraction curves for these parameter samples in panel (b) and the Ab-concentration dependent expected state durations in panel (c). We note that all parameter combinations in Θ_0.05,3_ had diffusing states that lasted less than 0.1 seconds for all values of [*A*]_exo_. We repeated this analysis for all donors and in each case found that 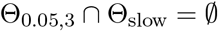.

### 3.3. Threshold and binding cascade parameters allow slow switching

By allowing the immobilization process to require multiple cross-linking antibodies, *T* > 1, and for the Ab-mucin dynamics to be state-dependent, *c* ≠ 1, we found both that (1) the subset of parameters that lead to slow switching is non-empty (Θ_slow_ ≠ 0), and (2) there is an overlap between slow-switching parameters and parameters that fit the Immobilized Fraction data well (Θ_0.05,5_ ∩ Θ_slow_ ≠ 0). For example, in Figure 7 panels (d)-(f) we demonstrate this fact assuming *T* = 19 for Donor F08. The 2*d* profile likelihood plot, in panel (d) shows an inverse relationship between *N*_*_ and the cascade factor *c*. Again the black region corresponds to all (*c, N*\*) pairs that appear in Θ_0.05,5_. For a. uniform sample of such pairs, in panel (e) we display the Immobilized Fraction predictions, and in panel (f) the corresponding expected immobilization and diffusion state durations. Only a. small subset, of Θ_0.05,5_ allows for slow switches. We mark this subset, in blue in all three panels. Notably, conditioned on *T* = 19, we have that *N*_*_ ≤ 120, which is somewhat smaller than the typical estimate for *N*_*_. In the next, section we note that assuming higher values for *T* leads to higher allowable values for *N*_*_. This type of result, holds for all donors: for sufficiently high assumed *T*, the corresponding parameters sets Θ_0.05,5_ and Θ_slow_ overlap.

By testing over a. range of 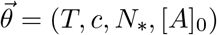, we uncovered some relationships among the components of the parameter vectors 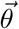 that yield slow switching Θ_slow_. We first, investigated the relationship between *T* and *c* by fixing *N*_*_ and [*A*]_0_. Noting that 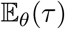 is independent, of *c* and 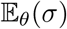 is an increasing function in *c*, we calculated the minimal *c* required to satisfy the slow switching condition, labeling this value *c*_min_. Though we could not. obtain an explicit, relationship between *T* and *c*_min_, we found that virions with a. large immobilization threshold *T* can only satisfy the slow switching condition if there is a. corresponding large cascade effect., large *c*_min_. To visualize this, in Figure 8(a) we display the parameter combinations of (*T,c, N*_*_ = 300, [*A*]_0_ = 0.1) that yield 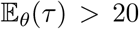 (green) and 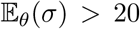 (red) for all exogenous antibody concentrations between 0 and l*μ*g/mL. The overlapping region (blue points) corresponds to *θ* ∈ Θ_slow_ and the combinations of interest (*T, c*_min_) are shown in black.

**Figure 8.**
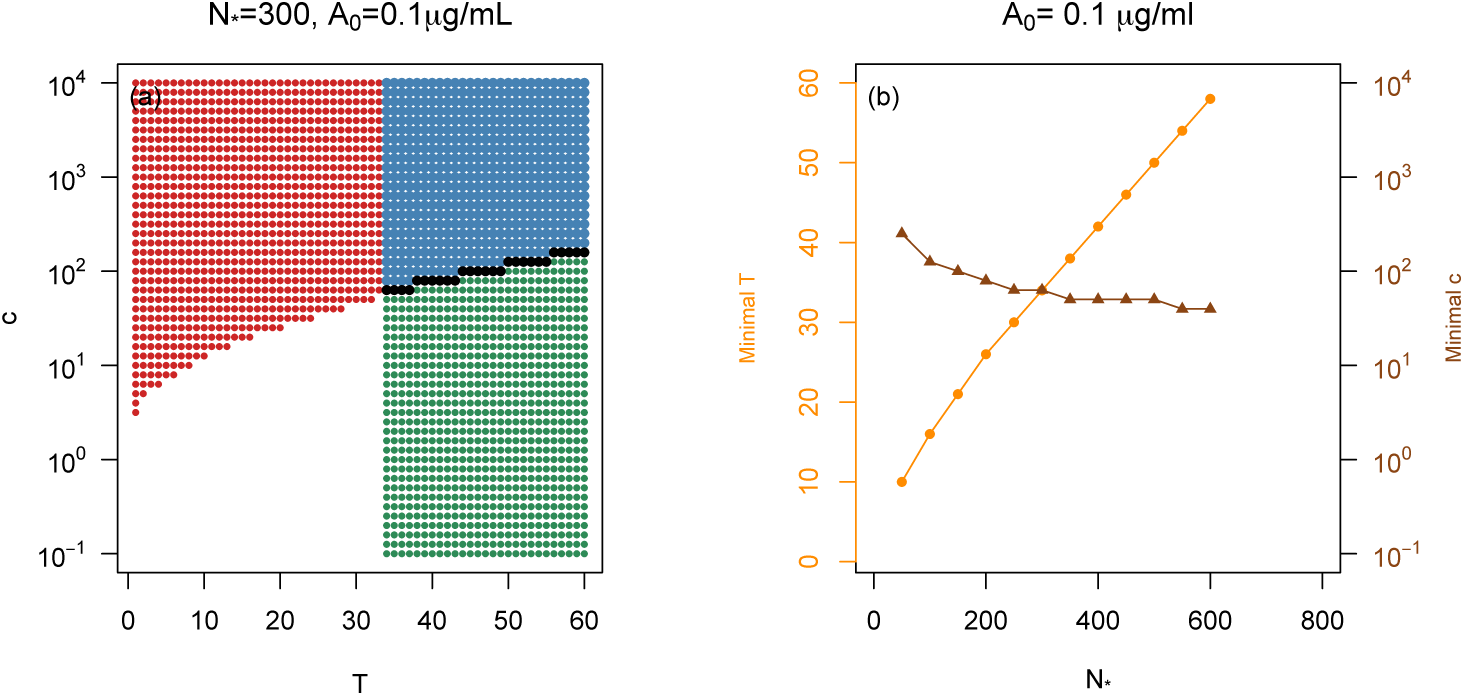
***(a)** Parameter combinations of T and c that predict expected immobilized times greater than 20s, red points, and predict expected freely-diffusing tim.es greater than 20s, green points, assuming N_*_ = 300 and [A]_0_ = 0.1 μg/m.L. The overlapping (T,c) combinations (blue points) are those combinations that satisfy slo w switching condition and subset (T, c_min_) are denoted by black, **(b)** The minimal value ofT required for our m.odel to predict slow switching as a function of N_*_, orange curve. Given an N_*_ and corresponding minimal T pair, the minimal value of c required for our m.odel to predict slo w switching is denoted by the brown curve. The endogenous Ab concentration is fixed at [A]_0_ = 0.1 μg/m.L*.

We draw the conclusion that if *N*_*_ = 300, then *T* must be at least 34 and *c* must be at least 63. If we increase the assumption about *N*_*_ while keeping [*A*]_0_ fixed, then we found that the minimal allowable *T* and *c* for slow switching increase and decrease, respectively. We demonstrate this relationship in Figure 8(b). For [*A*]_0_ = 0.1*μ*g/mL, the orange (circles) curve corresponds to the minimal *T* value (left y-axis) for the given *N*_*_ (x-axis) required such that 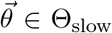 where [*A*]_0_ = 0.1*μ*g/mL. The brown (triangles) curve denotes the minimal *c* value (right *y* axis) required for the given *N*_*_, minimal *T*, and [*A*]_0_ = 0.1*μ*g/mL to result in expected state times longer than 20 seconds.

### 3.4. Model with threshold and binding cascade parameter is unidentifiable

As implied by the results in the preceding section, we found that the introduction of *T* > 1 and *c* ≠ 1 resulted in issues with identifiability. That is to say, it appears that the confidence region Θ_0.05,5_ is infinite even when restricted to the subspace Θ_0.05,5_∩Θ_slow_. We use the Immobilized Fraction data for Donor F08 to demonstrate this fact but provide information for each Donor in the Supplementary Information. Throughout this section we will use the terminology defined in subsection 2.6.

Over the full parameter space Θ, the Id profile likelihoods revealed that all three of the parameters *T, c*, and *N*_*_ are practically unidentifiable over the range we tested. The profile likelihoods are displayed in black in Figure 9(a)-(c). When we profiled the parameters *T, c*, and *N*_*_ restricted to the Slow Switching Regime Θ_0.05,5_ ∩ Θ_slow_, we found *T* is still practically unidentifiable over the range *T* ≥ 19, while c is practically unidentifiable a large range of positive values. The number of binding sites *N*_*_ does seem to be identifiable, with a deep valley centered around the unique minimum at approximately *N*_*_ = 120. These profile likelihoods are represented in blue in Figure 9(a)-(c). The dashed lines correspond to the 95% confidence interval boundaries for each parameter. Since the blue curves are below the confidence interval we can say that there exist parameter combinations in the Slow Switching Regime that reasonably fit the Immobilized Fraction data in Figure 9(e).

**Figure 9.**
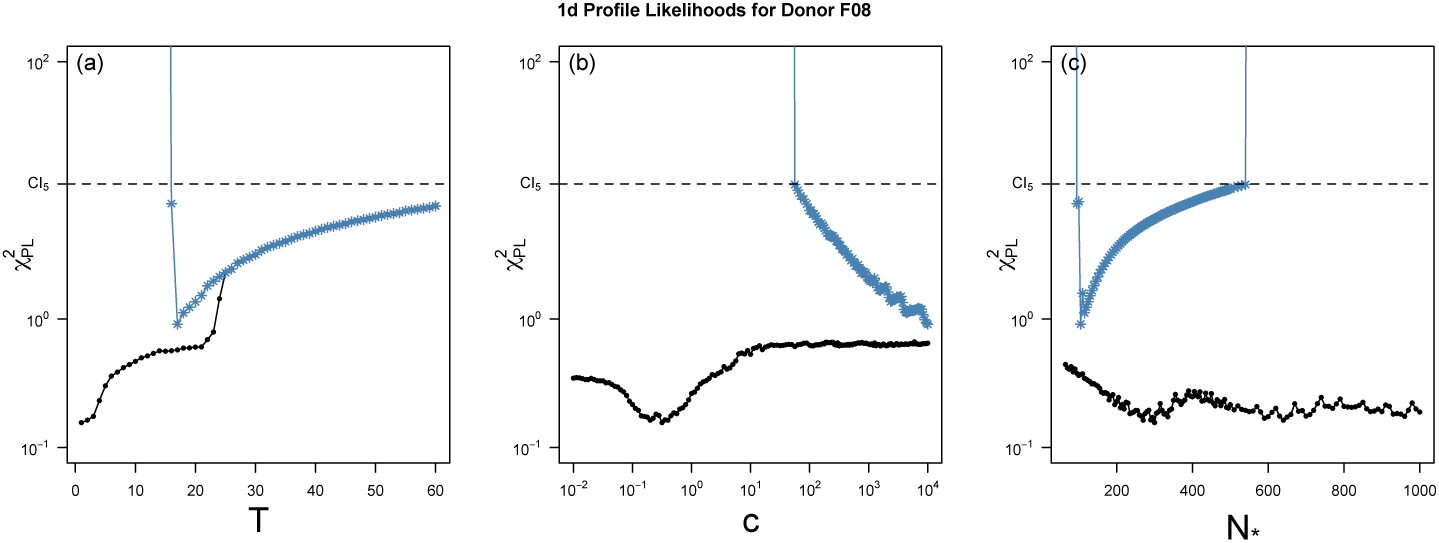
*(a)-(c) The Id profile likelihoods for the param.eters: immobilization threshold T, cascade factor c, and number of Ab binding sites on the virion N_*_, respectively over all tested param.eter combinations (black curves) and when restricted to the slow switching regim.e (blue curves). The 95% confidence interval for each param.eter consists of those param.eter values with, profile likelihood values below the dashed line*.

## 4. Discussion

We have developed mathematical models and statistical methods to analyze the behavior of HSV virions diffusing in CVM in the presence of various concentrations of cross-linking Ab. With a few exceptions, we found that particle paths can be partitioned into two basic categories: Freely Diffusing and Immobilized. While the fraction of Immobilized virions increases with Ab concentration, we found that the mobility of the Freely Diffusing class is not Ab-concentration dependent.

Because we expect all the individual bonds to be reversible, virions should switch between the Freely Diffusing and Immobilized states. Previously, it had been hypothesized that such switches are rapid with respect to the experimental time scale, but our analysis contradicts that assumption. This raises the question of whether or not it is possible for the basic kinetic model to produce “slow-switching” paths where switches occur on a time scale much larger than the experimental time window. We found that this is possible if the model allows for a lower bound on the number of Ab necessary to immobilize a virion and assuming a “cascade effect” in Ab-mucin binding that encourages entanglement.

Introducing these extra features leads to a fundamental issue with unidentifiability in the statistical analysis. We can make claims like “the minimum number of antibodies needed to immobilize a virion must be greater than 20 or so”, but we cannot be more specific. In order to do so, we would need to have access to time series that are much longer than what is currently experimentally feasible.

While we have shown that it is possible for reversible kinetics to be consistent with the path data, it might also be possible to explain the data with a model that assume all binding events are irreversible. Unfortunately the available data cannot distinguish between the two models. One possible resolution is to conduct experiments that explicitly control for the time between the introduction of Ab to the virion population and the observation of virion trajectories. Based on our model, in which we assume the immobilization process is reversible prior to the system reaching stationarity, switching should be more common when the number of antibodies bound to surface epitopes is low. Therefore, starting the tracking immediately enhances the probability of observing state switches before any long-lasting immobilization events occur.

On the other hand, observing virions at different time points long after Ab introduction will help determine whether or not the system reaches a stationary distribution. If so, there should be substantial information in analyzing how (or if) that stationary distribution depends on the Ab concentration, and the rate at which that stationary distribution is achieved.

The statistical methods and mathematical model introduced here apply to a broad class of biological systems that are composed of distinct subpopulations. Our classification scheme based on path-by-path analysis detects subpopulation dynamics that can be masked when considering only overall ensemble behavior. Clustering and then analyzing subpopulation ensemble statistics provides insight on the way the proportion and dynamics of these subpopulation change in response to the environmental factors. The model proposed in subsection 2.4, can be modified to describe the general scenario when nanoparticles work to entrap a diffusing pathogen by anchoring the pathogen to the surrounding environment.

## Supporting information

All supplementary materials

## Appendix A. Derivation of the MLE for D

From the defining properties of Brownian motion, the likelihood function of 2d Brownian motion defined by 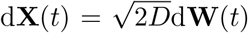 has form

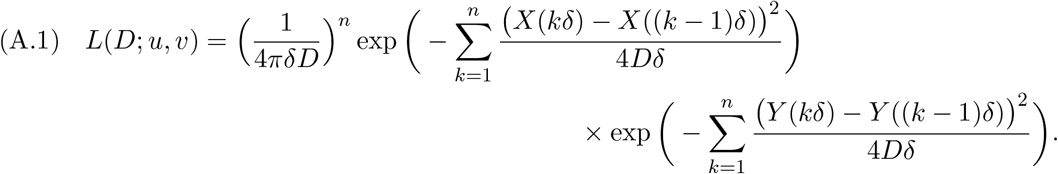

In terms of the increment process, *U*(*kδ*) and *V*(*kδ*), the loglikelihood is

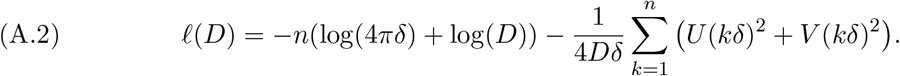

Solving the likelihood equation 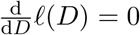, the ML estimator for *D* is given by

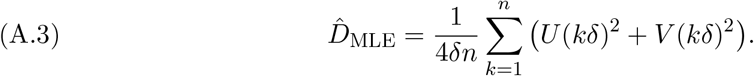

## Appendix B. Mathematical Model in subsection 2.4

We arrive at our approximation to the probability of immobilization Equation (2.10) presented in subsection 2.4 by averaging over the transitions of the number of antibodies simultaneously bound to the virion, *S*(*t*). To do this, we consider a simplified model in which the Ab-mucin binding rate is the same for both an immobilized virion and freely diffusing virion. That is the function defined in Equation (2.9) is constant, *g*(*s*) ≡ *c*. In this case, let *S*(*t*)|_*n,c*_ denote the Markov chain with transition rates:

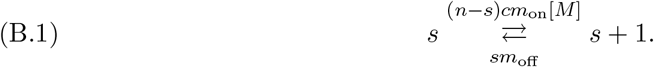

First, we derive the stationary distribution for the number of bound antibodies, *N*(*t*), and the conditional number of simultaneously bound antibodies assuming *g*(*s*) ≡ *c*, *S*(*t*)|_*n,c*_. Then we compute the expected duration of the Immobilized state and the Freely-Diffusing state of a virion from those quantities assuming *g*(*s*) ≡ *c*. Finally, we obtain Equation (2.10) using the results of the previous two steps.

### B.1. Stationary distribution of the two processes *N*(*t*) and *S*(*t*)|_*n;c*_

We model the two processes *N*(*t*) and *S*(*t*)|_*n;c*_ as CTMC with transition rates given by Equation (2.7) and Equation (B.1), respectively. Because they are irreducible Markov chains with a finite state space, there exists a unique stationary distribution, and convergence is exponential. Under the assumption that the Ab-binding sites on the surface of a virion operate independently, the process *N*(*t*) follows a binomial distribution with *N*_*_ trials and a time dependent success probability. Evoking a classical result from Renewal Theory, the steady state success probability is given by the long run fraction of being in the bound state, so that

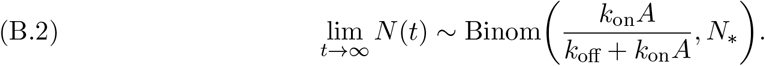

By assuming *g*(*s*) ≡ *c*, each antibody bound to the virion interacts with the mucin fibers independently, so that *S*(*t*)|_*n;c*_ is a binomial random variable with n trials and time dependent success probability. It follows from the same reasoning as above, that

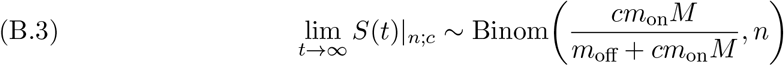

is the unique stationary distribution.

### B.2. Expected duration of the Freely-Diffusing and Immobilized states

We derive the expected duration of the Freely-Diffusing state, *τ*, and Immobilized states, *σ*, of a virion by considering the simplified model when the number of simultaneous bound antibodies has transition rates Equation (B.1).

We introduce the following notation *τ_T;c_* and *σ_T;c_* denote the expected time the process S_n;c_ spends in the freely diffusing state and immobilized state, respectively. The expected duration of the Freely-Diffusing state, *S_n;c_*(*t*) < *T*, is simply the expected hitting time of state *T*, given *S_n;c_* starts with *T* − 1 simultaneously bound antibodies. By solving a system of linear equations for the vector of expecting hitting times of state *T*, see [12], yields

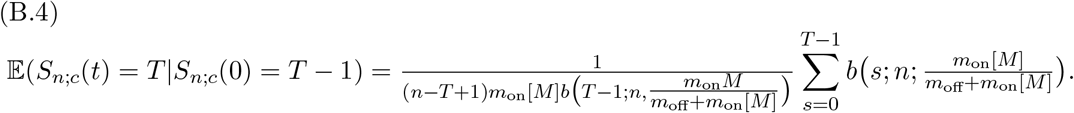

Similarly, the expected duration of the Immobilized state, *S_n;c_*(*t*) ≥ *T*, is the expected hitting time of state *T* − 1 given *S_n;c_* starts in state T. By solving a system of linear equations for the vector of expecting hitting times of state *T* − 1,

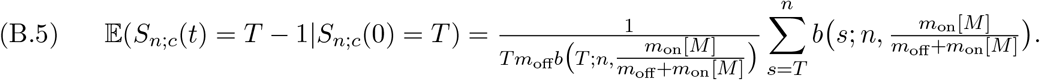

We observe that the transition rates of a Freely Diffusiving virion are the same as a virion modeled by the simplified transition rates given by Equation (B.1), specifically for *c* = 1. The transition rates of an immobilized virion are the same as a virion modeled by the simplified transition rates Equation (B.1). Hence, the following equalities hold

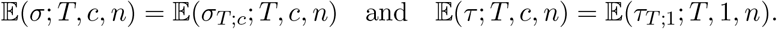

Explicitly, the duration of the Freely-diffusing state and the Immobilized state of a virion are given by

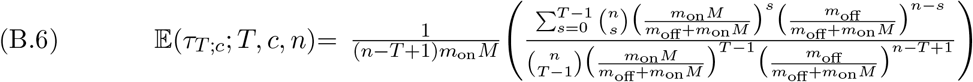

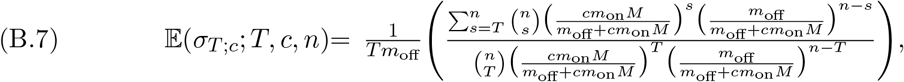

respectively.

### B.3. Asymptotic probability of number of bound antibodies

We are now ready to derive our time-scale approximation of the asymptotic probability of immobilization function, Equation (2.10). By conditioning on the slow process, the antibody-virion dynamics,

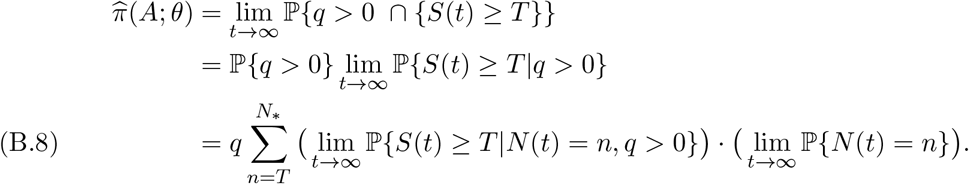

An application in Renewal Theory leads to the stationary probability of immobilization for the conditional process *S*(*t*) of the form

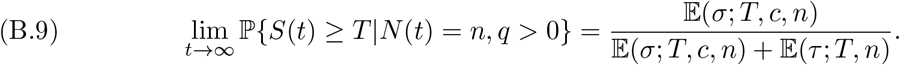

Plugging in the results from Appendix B.1 and Appendix B.2, gives Equation (2.10).

## Appendix C. Derivation of likelihoods

We derive the likelihood of Equation (2.14) and Equation (2.15) from the exact solution of the SDEs under the assumptions that the switch point, *τ*, occurs at a time measurement, and the true 2d position of the particle is **X**(*t*) = (*X*(*t*), *Y*(*t*)). We denote the time measurement *t_k_* = *kδ* for *k* = 1,… *n*, where *t*_0_ = 0 and *t_n_* = *T*, and **X**(*t_k_*) = **X**_*k*_.

For the [diffusion → immobilization] model, when *t* > *τ* the SDE is linear with additive noise. Hence a conditional exact solution can be expressed using Duhamel’s formula,

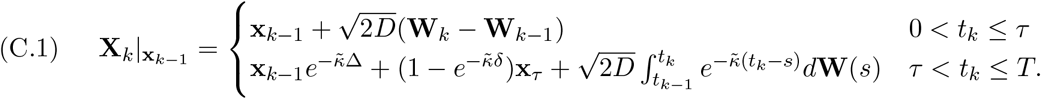

It follows immediately from the definition of Brownian motion and from an application of Ito’s Isometry,

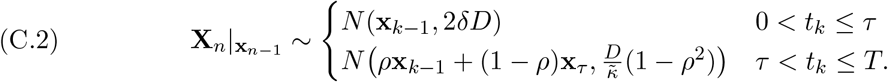

where 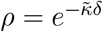. Because the solutions to SDEs satisfy the Markov Property,

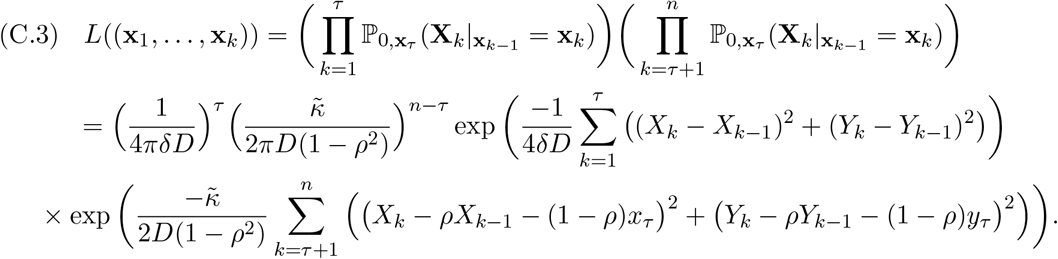

The likelihood equation for the [immobilization → diffusion] switching model derivation is similar to that [diffusion → immobilization] switching model but now we assume the immobilized particle is centered around the origin. Under the same reasoning as above

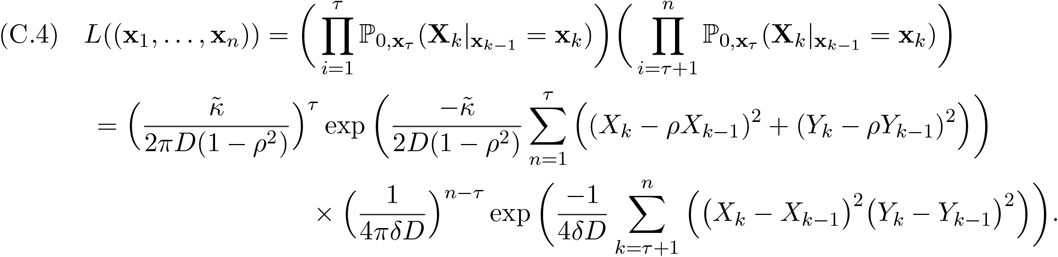

## Acknowledgments

The authors would like to thank Jay Newby, John Fricks, and Michelle Lacey for the helpful conversations that contributed to the development of this work.

