## Supplementary material for "Antibody-mediated immobilization of virions in mucus": All supplementary materials

**SM1. Typical virion trajectories.** We display the typical trajectories of the four different types of classified particles: Freely Diffusing, Immobilized, Subdiffusive, and Outlier in Figure SM1. The Freely Diffusing class made up 45.5246% of the paths (1689/3707), Immobilized class made up 53.0618% of the paths (1967/3707), Subdiffusive class made up 0.3507% of the paths (13/3707), and the outlier class made up 1.0251% of the paths (38/3707),

**SM2. Simulations of the immobilization process.** We simulate the immobilization process, depicted in Figure 3 using a Gillespie algorithm. First, we simulate the number of number of bound Ab to the surface of a virion, $N(t)$. To do this, we use the transition rates given in Equation (2.7) to randomly select if a bound Ab is gained or lost and randomly sample the time the virion remains in the current state.

Then for the given value of $N(t) = n$, we simulate the number of simultaneously bound Ab, $S(t)$ for the duration of time $N(t)$ remains in given state. Because we assume the Ab-mucin dynamics are fast compared to Ab-virion dynamics, we approximate $S(t)$ by a continuous space process using the stochastic differential equation

$$d\tilde{S}(t) = \big(\lambda(\tilde{S}(t)) - \mu(\tilde{S}(t))\big)dt + \sqrt{(\lambda(\tilde{S}(t)) + \mu(\tilde{S}(t)))}dW(t) \tag{SM2.1}$$

where $\lambda(\tilde{S}(t)) = \big(N(t) - \tilde{S}(t)\big)g(\tilde{S}(t))m_{\text{on}}[M]$ (the rate of gaining a simultaneously bound Ab) and $\mu(\tilde{S}(t)) = \tilde{S}(t)m_{\text{off}}$ (the rate of losing a simultaneously bound Ab). Then we use the Euler–Maruyama method to simulate the process $\tilde{S}(t)$ for the duration of time the number of bound Ab remains fixed.

**SM3. Clustering Algorithm Decision.** For each donor we give a table of the classification of each cluster at each tested exogenous antibody concentration and graphically display the grouping of clusters based on the path-wise statistics. We provide a description of each of the four subpopulations in Table SM1.

**SM3.1. Donor F02.** The dendrogram for Donor F02 was cut at a uniform height to yield four cluster for each tested exogenous Ab concentration except for $[A]_{\text{exo}} = 1.0\mu\text{g/mL}$, which was cut at a uniform height to yield five clusters because one particle was separated into its

*Submitted to the editors December 15th, 2018.

**Funding:** NIH R01GM122082-01, R21AI093242, U19AI096398, NSF DMS-1462992, DMS-1517274, CAREER Award DMR-1151477

†Department of Mathematics, Tulane University, New Orleans, LA.

‡Department of Biophysics, Johns Hopkins University, Baltimore, Maryland.

§Eshelman School of Pharmacy, University of North Carolina, Chapel Hill, Chapel Hill, NC.

¶Department of Mathematics, University of North Carolina, Chapel Hill, Chapel Hill, NC.

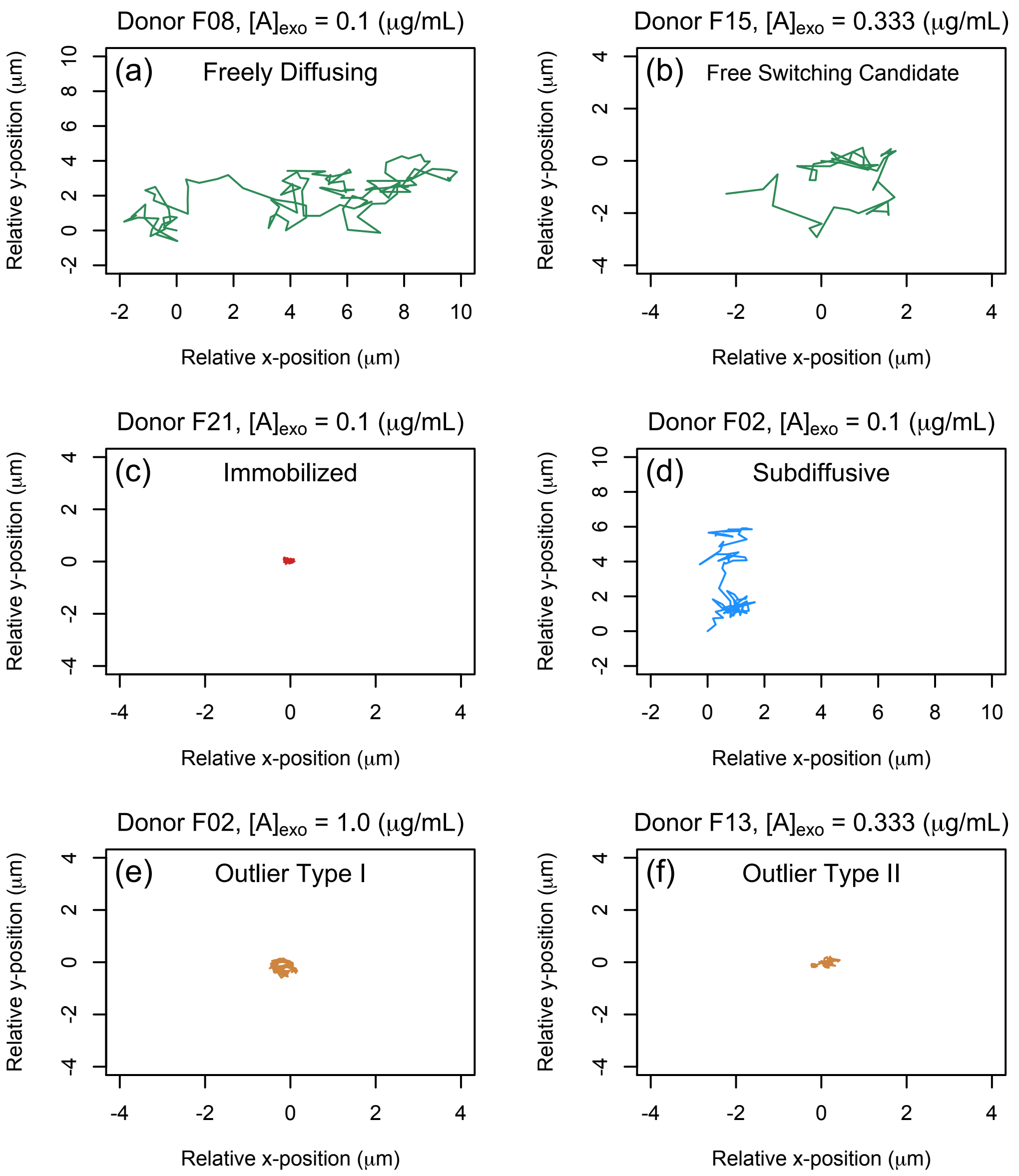


**Figure SM1.** *(a)-(f): A typical 2d-trajectory of a Virion classified as Freely Diffusing, Immobilized, Subdiffusive, Freely-Diffusing switch candidate, Outlier (Type I) and Outlier (Type 2), respectively.*

own cluster. The biological label that we assign to each cluster is reported in Table SM2. We display the clusters based on the path-wise statistics in Figure SM2, where the number corresponds to the hierarchical cluster and the color corresponds to the biological interpretation of the cluster.

**SM3.2. Donor F05.** The dendrogram for Donor F05 was cut at a uniform height to yield four cluster for each tested exogenous Ab concentration. The biological label that we assign

**Table SM1**

*Ensemble statistics for each of the four biological subpopulations.*

| | Diffusing | Immobilized | | Subdiffusive | Outlier |
|---|---|---|---|---|---|
| $\langle D_{\text{eff}} \rangle$ | $\geq 10^{-1}$ | $< 10^{-1}$ | | $\geq 10^{-1}$ | - |
| $\mathcal{A}(1;X), \mathcal{A}(1;Y)$ | Independent | Independent | Anti-persistant | Anti-persistant | - |
| Motion | Brownian | Hindered Brownian | Stationary | Subdiffusive | - |
| Color | Green | Red | | Blue | Brown |

**Table SM2**

*The classification of the clusters separated by the hierarchical clustering algorithm described in subsection 2.3. Short hand notation: $I$ = immobilized, $F$ = Freely Diffusing, $S$ = Subdiffusive, and $O$ = Outlier. We indicate a member of a cluster was removed and classified as Outlier by $-/O$.*

| $[A]_{\text{exo}}(\mu\text{g/mL})$ | Cluster 1 | Cluster 2 | Cluster 3 | Cluster 4 | Cluster 5 |
|---|---|---|---|---|---|
| 0 | I | I | F | F | - |
| 0.033 | I | I | I | F | - |
| 0.100 | I | I | F/O | F | - |
| 0.333 | I | I | I | F | - |
| 1.0 | I | O | S/O | S | F |

to each cluster is reported in Table SM3. We display the clusters based on the path-wise statistics in Figure SM3, where the number corresponds to the hierarchical cluster and the color corresponds to the biological interpretation of the cluster.

**Table SM3**

*The classification of the clusters separated by the hierarchical clustering algorithm described in Section 2.3. Short hand notation: $I$ = immobilized, $F$ = Freely Diffusing, $S$ = Subdiffusive, and $O$ = Outlier. We indicate a member of a cluster was removed and classified as Outlier by $-/O$.*

| $[A]_{\text{exo}}(\mu\text{g/mL})$ | Cluster 1 | Cluster 2 | Cluster 3 | Cluster 4 |
|---|---|---|---|---|
| 0 | I | I | F | O |
| 0.033 | I | I | S | F |
| 0.100 | I | I | I | F |
| 0.333 | I | I | I | F |
| 1.0 | I | I | I | F |

**SM3.3. Donor F08.** The dendrogram for Donor F08 was cut at a uniform height to yield four cluster for each tested exogenous Ab concentration. The biological label that we assign to each cluster is reported in Table SM4. We display the clusters based on the path-wise statistics in Figure SM4, where the number corresponds to the hierarchical cluster and the color corresponds to the biological interpretation of the cluster.

**SM3.4. Donor F13.** The dendrogram for Donor F13 was cut at a uniform height to yield four cluster for each tested exogenous Ab concentration except for $[A]_{\text{exo}} = 1.0\mu\text{g/mL}$, which was cut at a uniform height to yield five clusters because one particle was separated into its

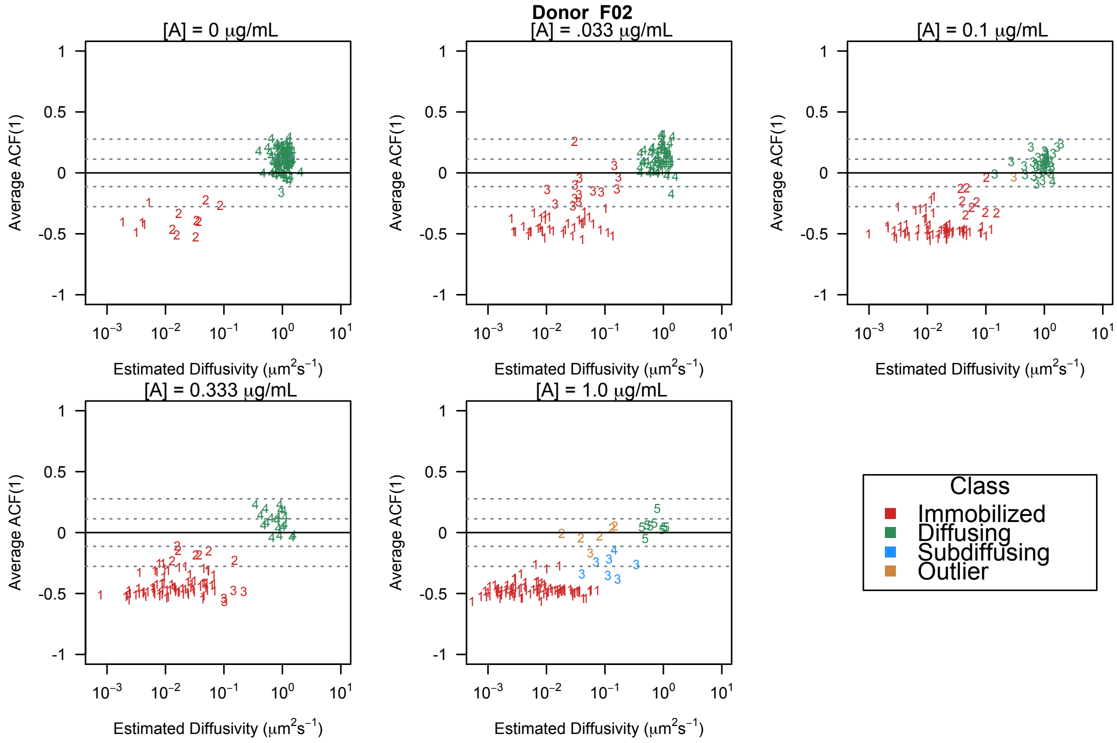


**Figure SM2.** *(a)-(e)A depiction of the classification system, where each point, corresponds to a tracked virion with estimated diffusivity on a* $\log 10$ *scale and average-ACF value, for Ab concentration* 0, 0.033, 0.1, 0.333, *and* $1.0\mu g/mL$*, respectively. The numerical value of each point corresponds to the prescribed cluster by the hierarchical clustering algorithm and the color of the point represents the biological class.*

**Table SM4**

*The classification of the clusters separated by the hierarchical clustering algorithm described in Section 2.3.. Short hand notation: I = immobilized, F = Freely Diffusing, S = Subdiffusive, and O = Outlier. We indicate a member of a cluster was removed and classified as Outlier by* $-/O$.

| $[A]_{\text{exo}}(\mu g/\text{mL})$ | Cluster 1 | Cluster 2 | Cluster 3 | Cluster 4 |
|---|---|---|---|---|
| 0 | I | F | F | F |
| 0.033 | I | I | F | F |
| 0.100 | I | I | S | F |
| 0.333 | I | I | F | F |
| 1.0 | I | I | S | F |

own cluster. The biological label that we assign to each cluster is reported in Table SM5. We display the clusters based on the path-wise statistics in Figure SM5, where the number corresponds to the hierarchical cluster and the color corresponds to the biological interpretation of the cluster.

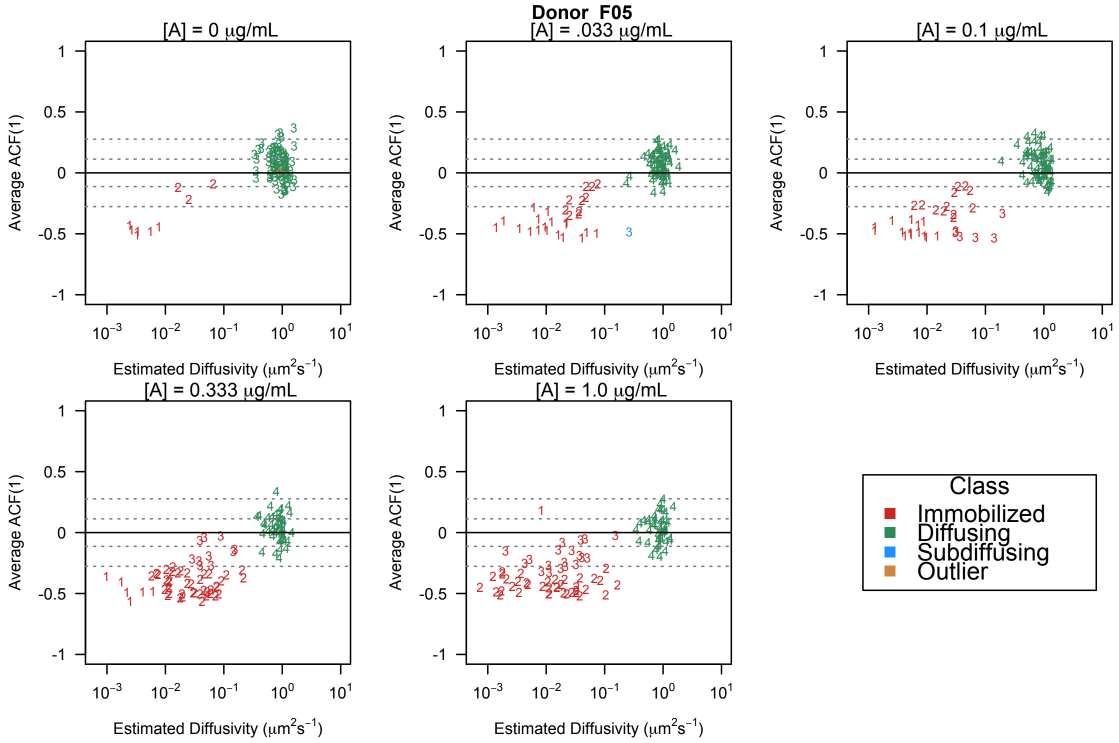


**Figure SM3.** *(a)-(e)A depiction of the classification system, where each point, corresponds to a tracked virion with estimated diffusivity on a* $\log 10$ *scale and average-ACF value, for Ab concentration* 0, 0.033, 0.1, 0.333, *and* $1.0\mu g/mL$*, respectively. The numerical value of each point corresponds to the prescribed cluster by the hierarchical clustering algorithm and the color of the point represents the biological class.*

**Figure SM4.** *(a)-(e)A depiction of the classification system, where each point, corresponds to a tracked virion with estimated diffusivity on a* $\log 10$ *scale and average-ACF value, for Ab concentration* 0, 0.033, 0.1, 0.333, *and* $1.0\mu g/mL$*, respectively. The numerical value of each point corresponds to the prescribed cluster by the hierarchical clustering algorithm and the color of the point represents the biological class.*

**SM3.5. Donor F15.** The dendrogram for Donor F15 was cut at a uniform height to yield four cluster for each tested exogenous Ab concentration. The biological label that we assign to each cluster is reported in Table SM6. We display the clusters based on the path-wise statistics in Figure SM6, where the number corresponds to the hierarchical cluster and the color corresponds to the biological interpretation of the cluster.

**SM3.6. Donor F17.** The dendrogram for Donor F17 was cut at a uniform height to yield four cluster for each tested exogenous Ab concentration. The biological label that we assign to each cluster is reported in Table SM7. We display the clusters based on the path-wise statistics in Figure SM7, where the number corresponds to the hierarchical cluster and the color corresponds to the biological interpretation of the cluster.

**Table SM5**

*The classification of the clusters separated by the hierarchical clustering algorithm described in Section 2.3. Short hand notation: I = immobilized, F = Freely Diffusing, S = Subdiffusive, and O = Outlier. We indicate a member of a cluster was removed and classified as Outlier by −/O.*

| $[A]_{\text{exo}}(\mu g/mL)$ | Cluster 1 | Cluster 2 | Cluster 3 | Cluster 4 | Cluster 5 |
|---|---|---|---|---|---|
| 0 | I | I | S | F | F |
| 0.033 | I | I | F | F | F |
| 0.100 | I | I | O | S | F |
| 0.333 | I | I | I | F/O | F |
| 1.0 | I | I | I | F | F |

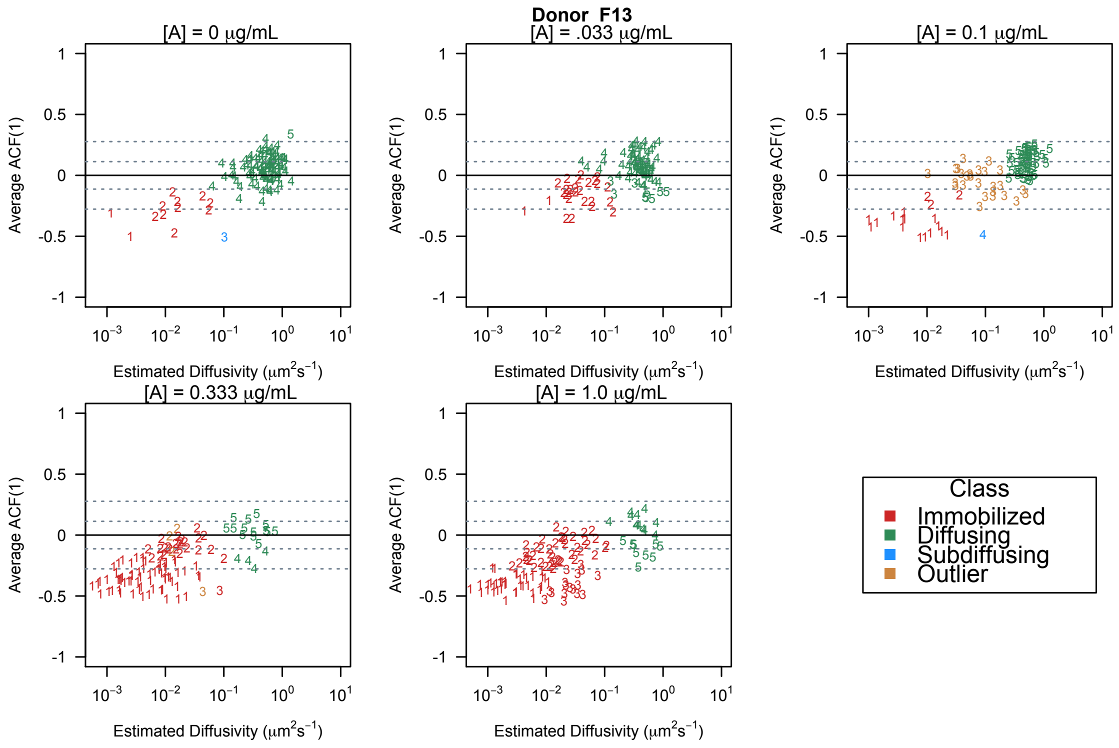


**Figure SM5.** *(a)-(e)A depiction of the classification system, where each point, corresponds to a tracked virion with estimated diffusivity on a* $\log 10$ *scale and average-ACF value, for Ab concentration* 0*,* 0.033*,* 0.1*,* 0.333*, and* $1.0\mu g/mL$*, respectively. The numerical value of each point corresponds to the prescribed cluster by the hierarchical clustering algorithm and the color of the point represents the biological class.*

**SM3.7. Donor F21.** The dendrogram for Donor 21 was cut at a uniform height to yield four cluster for each tested exogenous Ab concentration. The biological label that we assign to each cluster is reported in Table SM8. We display the clusters based on the path-wise statistics in Figure SM8, where the number corresponds to the hierarchical cluster and the color corresponds to the biological interpretation of the cluster.

**Table SM6**

*The classification of the clusters separated by the hierarchical clustering algorithm described in Section 2.3. Short hand notation: I = immobilized, F = Freely Diffusing, S = Subdiffusive, and O = Outlier. We indicate a member of a cluster was removed and classified as Outlier by* $-/O$.

| $[A]_{\text{exo}}(\mu\text{g/mL})$ | Cluster 1 | Cluster 2 | Cluster 3 | Cluster 4 |
|---|---|---|---|---|
| 0 | I | I | I | F |
| 0.033 | I | I | I | F |
| 0.100 | I | I | F | F |
| 0.333 | I | I | F | F |
| 1.0 | I | I | I | F |

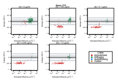

**Figure SM6.** *(a)-(e)A depiction of the classification system, where each point, corresponds to a tracked virion with estimated diffusivity on a* $\log 10$ *scale and average-ACF value, for Ab concentration* 0, 0.033, 0.1, 0.333, *and* $1.0\mu g/mL$, *respectively. The numerical value of each point corresponds to the prescribed cluster by the hierarchical clustering algorithm and the color of the point represents the biological class.*

### SM4. Switch Point Detection Algorithm.

**SM4.1. Hamiltonian Monte Carlo (HMC) Sampling Algorithm.** We used an HMC sampling algorithm to jointly estimate the parameters $(D, \tilde{\kappa}, \tau)$. Hamiltonian Monte Carlo is a form of Markov Chain Monte Carlo sampling that relies on Bayes Rule to sample from the joint posterior distribution of $(D, \tilde{\kappa}, \tau)$,

$$p(D, \tilde{\kappa}, \tau | \mathbf{x}, \mathbf{y}) \overset{c}{=} L(\mathbf{x}, \mathbf{y}; D, \tilde{\kappa}, \tau) p(D, \tilde{\kappa}, \tau) \quad \text{(SM4.1)}$$

where $L(\mathbf{x}, \mathbf{y}; (D, \tilde{\kappa}, \tau))$ is the likelihood of seeing the data given the parameters $(D, \tilde{\kappa}, \tau)$ and $p(D, \tilde{\kappa}, \tau)$ is the prior joint distribution of $(D, \tilde{\kappa}, \tau)$. In our model, we assume the parameters $D, \tilde{\kappa}$, and $\tau$ are independent so that the joint prior distribution factors into the product of priors, $p(D, \tilde{\kappa}, \tau) = p(D)p(\tilde{\kappa})p(\tau)$.

The likelihood function, $L(\mathbf{x}, \mathbf{y}; D, \tilde{\kappa}, \tau)$ is a consequence of the model. For the [diffusing $\rightarrow$ immobilized] switching scenario and [immobilized $\rightarrow$ diffusing] switching scenario $L(\mathbf{x}, \mathbf{y}; (D, \tilde{\kappa}, \tau))$ is given by Equation (2.15) and Equation (2.14), respectively.

As for the priors, we placed a loosely informative prior on $D$

$$\pi(D) \sim \text{Gamma}(D_{\text{eff}} * 0.1, 0.1)$$

and chose a discrete uniform distribution over the observation times $\{t_n\}_{n=1}^{N-1}$ for the prior distribution on $\tau$. Rather than directly sampling the parameter $\tilde{\kappa}$ , we sampled the transform $\rho = \exp(-\Delta\tilde{\kappa})$ and then used the inverse transformation $\tilde{\kappa} = -\log(\rho)/\Delta$ to obtain posterior samples for $\tilde{\kappa}$. In doing this, we expressed our uncertainty of the order of $\tilde{\kappa}$ by placing a lognormal prior on $\rho$,

$$\pi(\rho) \sim \text{lognormal}\Big(loc = \log\big(\tfrac{0.5^2}{\sqrt{0.5^2+2}}\big), shape = \log(1 + \tfrac{2}{0.5^2})^{1/2}\Big).$$

**Table SM7**

*The classification of the clusters separated by the ierarchical clustering algorithm described in Section 2.3.. Short hand notation: I = immobilized, F = Freely Diffusing, S = Subdiffusive, and O = Outlier. We indicate a member of a cluster was removed and classified as Outlier by −/O.*

| $[A]_{\text{exo}}(\mu\text{g/mL})$ | Cluster 1 | Cluster 2 | Cluster 3 | Cluster 4 |
|---|---|---|---|---|
| 0 | I | I | F | F |
| 0.033 | I | I | I | F |
| 0.100 | I | I | F | F |
| 0.333 | I | I | I | F |
| 1.0 | I | I | I | F |

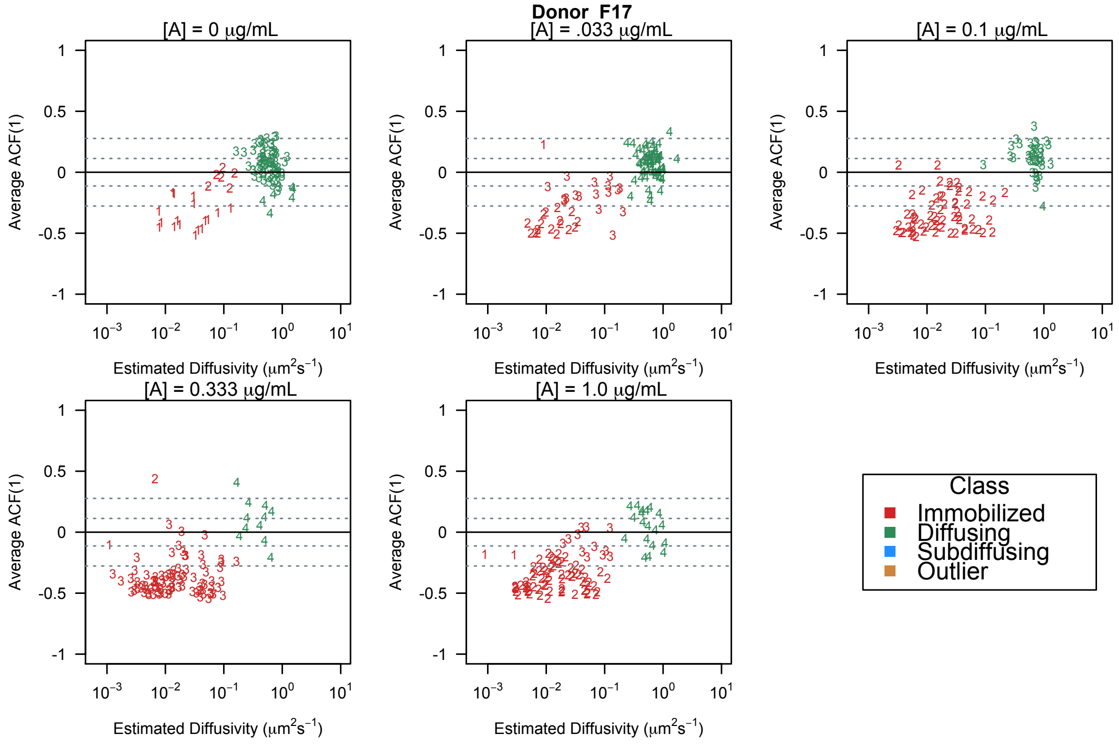


**Figure SM7.** *(a)-(e)A depiction of the classification system, where each point, corresponds to a tracked virion with estimated diffusivity on a* $\log 10$ *scale and average-ACF value, for Ab concentration 0, 0.033, 0.1, 0.333, and 1.0µg/mL, respectively. The numerical value of each point corresponds to the prescribed cluster by the hierarchical clustering algorithm and the color of the point represents the biological class.*

We note that because $\tau$ is a discrete parameter taking values in $\{t_n\}_{n=1}^{N-1}$ whereas $D, \tilde{\kappa}$ are continuous real-valued parameters, posterior samples of $\tau | D, \tilde{\kappa}$ were first drawn from $\{t_n\}_{n=1}^{N-1}$ with probability

$$p(\tau = t_j | \mathbf{x}, \mathbf{y}, D, \tilde{\kappa}) = \frac{L(\mathbf{x}, \mathbf{y}; D, \tilde{\kappa}, \tau = t_j) p(\tau = t_j)}{\sum_{k=1}^{N} L(\mathbf{x}, \mathbf{y}; D, \tilde{\kappa}, \tau = t_k) p(\tau = t_k)} \quad \text{for } j = 1, \ldots N-1. \tag{SM4.2}$$

**Table SM8**

*The classification of the clusters separated by the hierarchical clustering algorithm described in Section 2.3. Short hand notation: $I$ = immobilized, $F$ = Freely Diffusing, $S$ = Subdiffusive, and $O$ = Outlier. We indicate a member of a cluster was removed and classified as Outlier by $-/O$.*

| $[A]_{\text{exo}}(\mu\text{g/mL})$ | Cluster 1 | Cluster 2 | Cluster 3 | Cluster 4 |
|---|---|---|---|---|
| 0 | I | I | F | F |
| 0.033 | I | I | F | F |
| 0.100 | I | I | I | F |
| 0.333 | I | I | F | F |
| 1.0 | I | I | I | F |

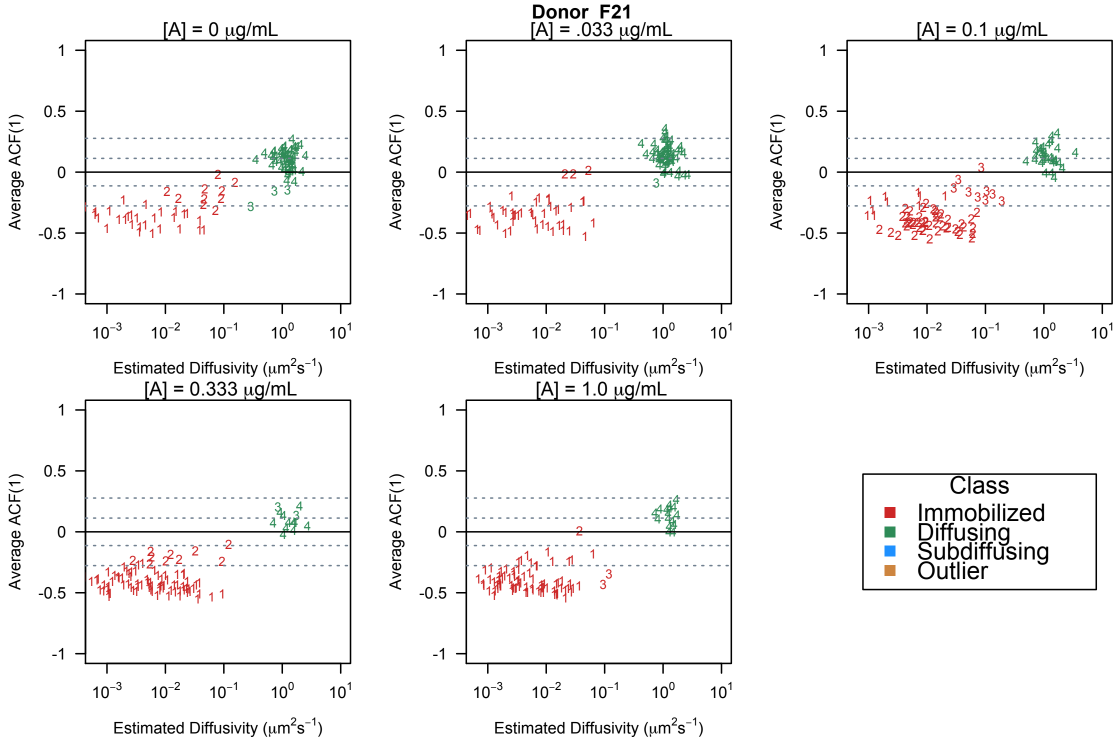


**Figure SM8.** *(a)-(e)A depiction of the classification system, where each point, corresponds to a tracked virion with estimated diffusivity on a $\log 10$ scale and average-ACF value, for Ab concentration 0, 0.033, 0.1, 0.333, and 1.0µg/mL, respectively. The numerical value of each point corresponds to the prescribed cluster by the hierarchical clustering algorithm and the color of the point represents the biological class.*

Then, we sampled $D$ and $\tilde{\kappa}$ using the marginal posterior distribution of $D$ and $\tilde{\kappa}$,

$$p(D, \tilde{\kappa}|\mathbf{x}, \mathbf{y}) \stackrel{c}{=} p(D)p(\tilde{\kappa}) \sum_{k=1}^{N} L(\mathbf{x}, \mathbf{y}; D, \tilde{\kappa}, \tau = t_k)p(\tau = t_k). \tag{SM4.3}$$

The HMC sampling algorithm was run using the `R` package rstan, and computations were done using high performance computing (HPC) resources and services provided by Technology Services at Tulane University, New Orleans, LA.

**SM4.2. Testing Freely Diffusing population for switch candidates.** Recall, we say a path is a candidate for switching if the 95% credible region for $\tau|D,\tilde{\kappa}$ is completely contained within the interval $[0.1\mathrm{T}_{\mathrm{final}},\ 0.9\mathrm{T}_{\mathrm{final}}]$ where $\mathrm{T}_{\mathrm{final}}$ is the duration of a path. We tested if the $i$-th Freely Diffusing virion of path duration $\mathrm{T}_{\mathrm{final}}$, was a [immobilization $\rightarrow$ diffusion] switching candidate using the following procedure. Assuming the [immobilization $\rightarrow$ diffusion] switching model, we obtained posterior samples of $\tau|D,\tilde{\kappa}$ using our HMC sampling algorithm, in which we drew 2000 posterior samples and kept the last 1000 samples for 4 different runs of the algorithm. The resulting 4000 posterior samples of $\tau|_{D,\tilde{\kappa}}$ were used to construct the 95% credible interval for $\tau$. If this interval was contained in $[0.1\mathrm{T}_{\mathrm{final}},\ 0.9\mathrm{T}_{\mathrm{final}}]$, then we marked the virion as a candidate for [immobilization $\rightarrow$ diffusion] switching. We repeated this for all virions in the Freely Diffusing diffusing subpopulation to obtain the fraction of candidate for [immobilization $\rightarrow$ diffusion] switching. We obtained the fraction of candidate for [diffusion $\rightarrow$ immobilization] switching, following the same method described in the paragraph above but assuming the [diffusion $\rightarrow$ immobilization] switching model.

The Gaussian kernel density estimate of the posterior samples of $\tau$ within the 95% credible region for [immobilization $\rightarrow$ diffusion] switch candidates, and [diffusion $\rightarrow$ immobilization] switch candidates are displayed in top Figure SM9 (a) and (b), respectively, for Donor F15. The path duration for each switch candidate is marked by a red $x$. In Figure SM9 (c) and (d), we display the trajectory of an [immobilization $\rightarrow$ diffusion] switch candidates and [diffusion $\rightarrow$ immobilization] switch candidate, where the start of the trajectory by the green dot and the maximum a posteriori estimate for $\tau$ is marked by the blue star.

**SM4.3. Estimating the false discovery rate.** To estimate the false discovery rate of our switch detection test, we simulated Feely diffusing trajectories using parameters obtained from the class of Freely Diffusing virions in our data. To do this, the number of simulated Brownian paths was set to the number of virions classified as Freely diffusing, 1689 virions. Let $\mathbf{n}_{\mathrm{free}}$ and $\mathbf{D}_{\mathrm{free}}$ denote all the path lengths and the effective diffusivities of the Freely Diffusing virions, respectively. For the $i$-th simulated Brownian path, the path length $n_i$ was sampled from $\mathbf{n}_{\mathrm{free}}$ and the diffusivity constant $D_i$ was sampled from $\mathbf{D}_{\mathrm{free}}$.

Using our switch point criterion, we set the false discovery rate of [Immobilization $\rightarrow$ Diffusion] switches to the percent of simulated Brownian particles that were labeled as candidates for [Immobilization $\rightarrow$ Diffusion] switching for a likelihood function given by Equation (2.15). Specifically, the $i$-th simulated Brownian path and [Immobilization $\rightarrow$ Diffusion] switching model, the 95% credible region for $\tau$ was constructed using the 4000 posterior samples of $\tau|_{D,\tilde{\kappa}}$ obtained by the last 1000 (out of 2000) posterior samples of 4 runs of our HMC algorithm. We obtained an estimate for the false discovery rate of [Diffusion $\rightarrow$ Immobilization] switches in the same manner but assuming a [Diffusion $\rightarrow$ Immobilization] model.

**SM4.4. Estimating the power of the candidate switching test.** In estimating the power of our test, we assumed that the switch point occurs within the middle 80% of the path. We believe this was a reasonable assumption because on average a Freely Diffusing virion had a path length of 80, and the first or last eight observations would be too noisy to capture the

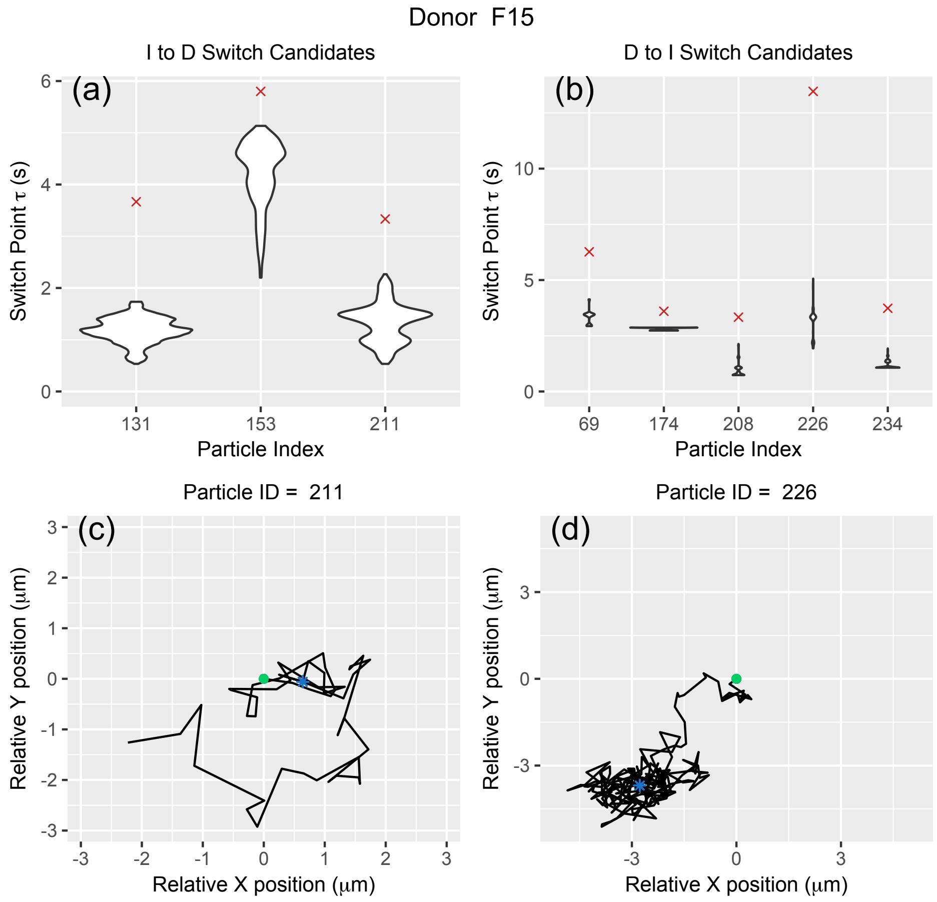


**Figure SM9.** ***(a)-(b)*** *Violin plots for the posterior samples of $\tau$ within the 95% credible interval for those Freely-Diffusing virions from Donor F15 considered [immobilization $\to$ diffusion] switch candidates, and [Diffusion $\to$ Immobilization] switch candidate, respectively.* ***(c)*** *The 2d trajectory [immobilization $\to$ diffusion] switch candidate, (tracked particle 211 for Donor F15 at $[A]_{exo} = 0.333\mu g/mL$.)* ***(d)*** *The 2d trajectory [diffusion $\to$ immobilization] switch candidate, (tracked particle 226 for Donor F15 at $[A]_{exo} = 1.0\mu g/mL$.) In Panels (c) and (d) the green circle denotes the initial position of the virion and the blue star corresponds to the estimated switch point.*

true dynamics of the virion. First, we created a simulated data set of 1689 [immobilization $\to$ diffusion] paths from the explicit solution of SDE given in Equation (2.14). For the $i$th [immobilization $\to$ diffusion] path, the model parameters $(n_i, D_i, \tilde{\kappa}_i, \tau_i)$ were sampled from the summary statistics of the tracked virions:

$$n_i \sim \text{Unif}\{\mathbf{n}_{\text{free}}\}, \quad D_i \sim \text{Unif}\{\mathbf{D}_{\text{free}}\}, \quad \tilde{\kappa}_i \sim \text{Exp}(\lambda = 1/500), \text{ and } \tau_i \sim \text{Unif}\{[0.1n_i, 0.9n_i]\}.$$

We ran our sampling algorithm assuming the model given by Equation (2.15) on the simulated [immobilization $\to$ diffusion] data. The sampling algorithm was run 4 times each for 2000 iterations where the last 1000 were kept as posterior samples. We then set the power of our [immobilization $\to$ diffusion] switch point detecting test to the percentage of them have credible regions for $\tau$ within $[0.1T_{\text{final}}, 0.9T_{\text{final}}]$. The test failed to detect switches in the following two cases: (1) The switch was located near the endpoints of the interval $[0.1T_{\text{final}}, 0.9T_{\text{final}}]$, resulting in a credible region outside the region allowed by the test. (2) When $\tilde{\kappa}$ is approximately less than 10. In this case, the immobilized state was indistinguishable from Brownian Motion because the deterministic component of the SDE given in Equation (2.15) was relatively small with respect to the random component. In Figure SM10(a)-(b), we display the relative position of the change point and $\tilde{\kappa}$ for simulated paths that were undetected by our proposed test. The trajectory of the simulated path marked by the pink diamond is shown in Figure SM10(c)-(d).

We estimated the power of [diffusion $\to$ immobilization] switch point detecting test following the same procedure as above but assumed the model given by Equation (2.14). For this switching scenario, switches went undetected due to the same reasons mentioned above.

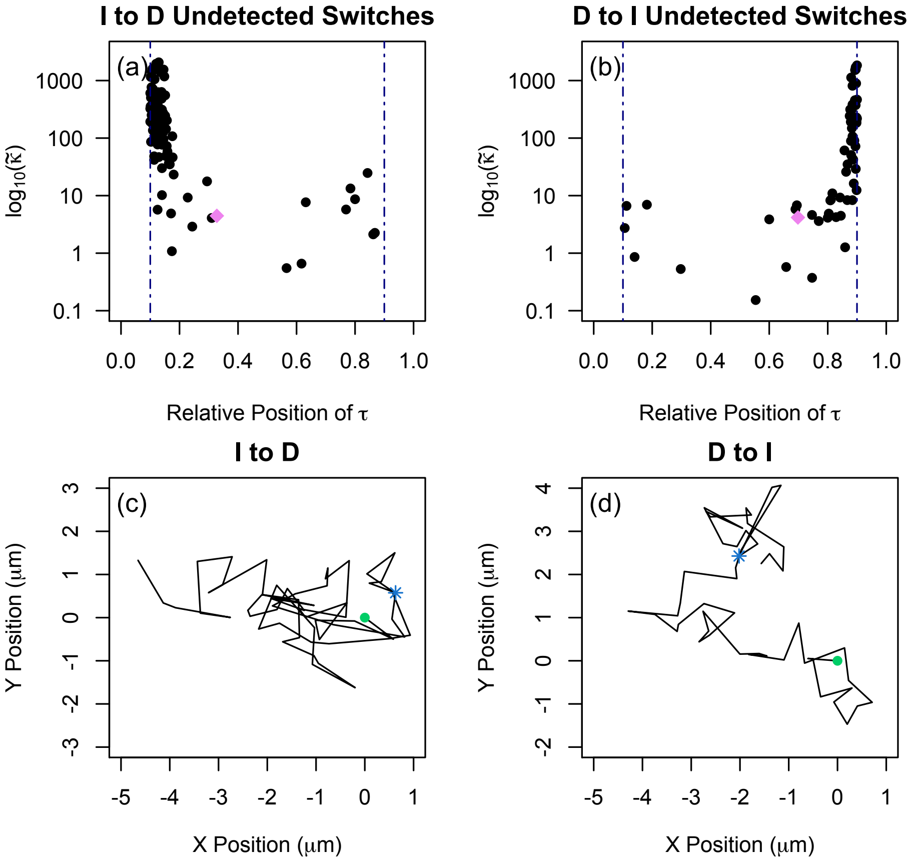


**Figure SM10.** ***(a)-(b)*** *The true parameter $\tilde{\kappa}$ and the relative position of the switch point, $\tau$, for the simulated switching particles, [immobilization $\to$ diffusion] and [diffusion $\to$ immobilized], respectively, whose switch point was undetected by our switching detection test. The dashed navy vertical lines mark the tenth percentile and ninetieth percentile of the trajectory.* ***(c)-(d)*** *The trajectory of a simulated switching particles marked by the pink diamond panels (a) and (b), respectively.*

**SM5. Statistical Evidence for trend in proportion immobilized.** In Figure SM11, we display the 95% $BC_a$ confidence interval. for the observed proportion immobilized for each donor. We remark that for all donors, the proportion immobilized at the lowest tested exogenous Ab concentration ($0\mu$g/mL) and the highest tested exogenous Ab concentration ($1\mu$g/mL) have non-overlapping confidence intervals. In Table SM9, we report the estimates coefficient and associated $p$-values for a negative exponential growth for the observed proportion of virions immobilized.

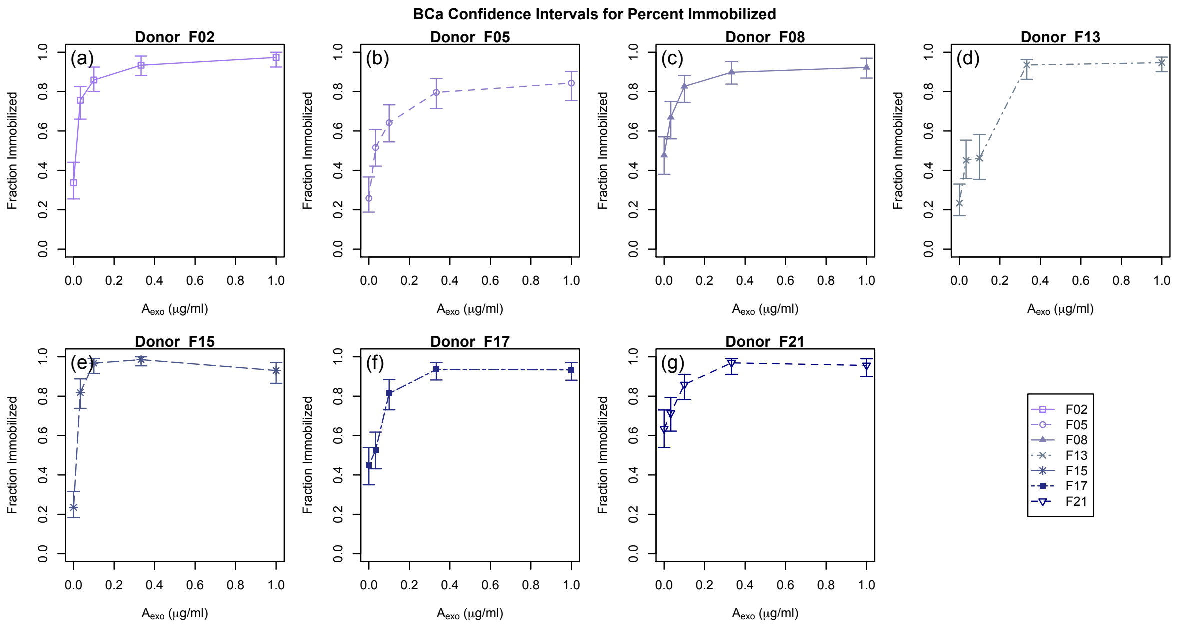


**Figure SM11.** *(a)-(g) For each donor, the observed proportion immobilized with 95% $BC_a$ confidence interval.*

**Table SM9**

*Negative exponential growth coefficient estimates and associated p-values for the observed fraction of virions immobilized.*

| | Estimate | $p$ -value |
|---|---|---|
| $\beta_0$ | 0.9138 | $< 0.0001$ |
| $\alpha_0$ | -0.8427 | 0.0043 |
| $\alpha_{\text{exo}}$ | 15.9201 | $< 0.0001$ |

| | Estimate | $p$ -value |
|---|---|---|
| $\beta_1$ | 0.0642 | 0.4547 |
| $\beta_2$ | -0.1084 | 0.2123 |
| $\beta_4$ | -0.0700 | 0.4156 |
| $\beta_5$ | 0.1140 | 0.1913 |
| $\beta_6$ | 0.0050 | 0.9534 |
| $\beta_7$ | 0.0347 | 0.6844 |

| | Estimate | $p$ -value |
|---|---|---|
| $\alpha_1$ | 0.2846 | 0.3952 |
| $\alpha_2$ | 0.2272 | 0.5049 |
| $\alpha_4$ | 0.4286 | 0.1852 |
| $\alpha_5$ | 0.4385 | 0.1745 |
| $\alpha_6$ | 0.1901 | 0.5819 |
| $\alpha_7$ | -0.2406 | 0.5763 |

**SM6. Statistical Evidence for Effective Diffusivity.** In Figure SM12, we display the 95% $BC_a$ confidence interval for the ensemble effective diffusivity for the Freely-Diffusing subpop-

ulation. We remark that for all donors, the ensemble effective diffusivity at the lowest tested exogenous Ab concentration (0$\mu$g/mL) and the highest tested exogenous Ab concentration (1$\mu$g/mL) have overlapping confidence intervals. We report the results of all the paired difference tests of the form,

$$H_0 : \langle D_{\text{eff}}([A]_i)\rangle - \langle D_{\text{eff}}([A]_j)\rangle = 0, \qquad H_A : \langle D_{\text{eff}}([A]_i)\rangle - \langle D_{\text{eff}}([A]_j)\rangle > 0. \tag{SM6.1}$$

for $1 \leq i < j \leq 5$ in Table SM10.

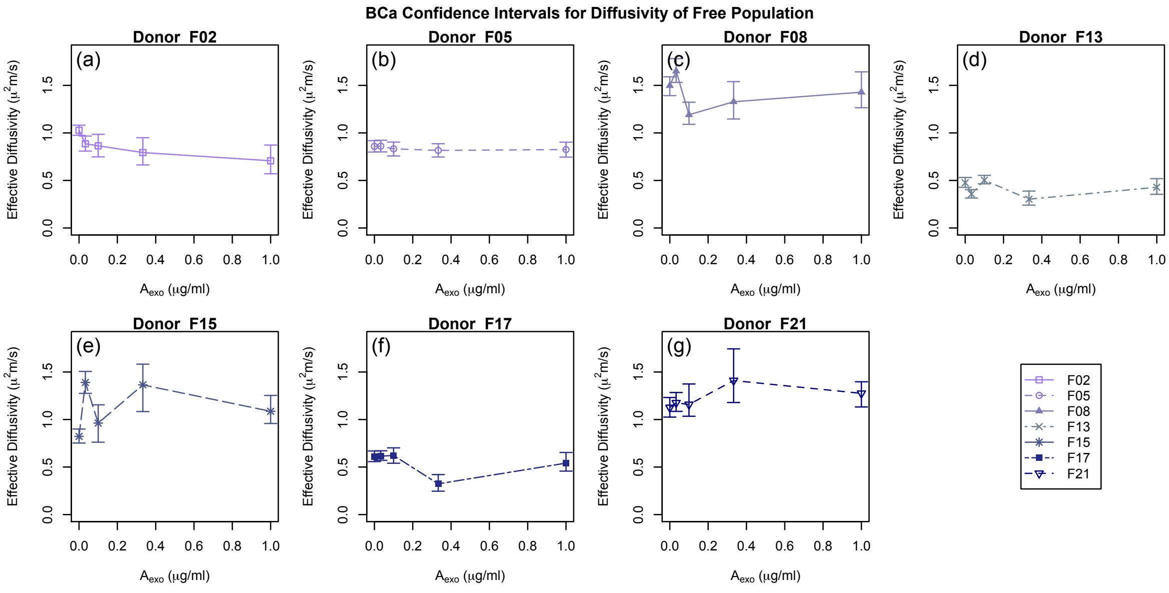


**Figure SM12.** *(a)-(g) For each donor, the ensemble effective diffusivity of the free subpopulation with 95% $BC_a$ confidence interval.*

**Table SM10**

*The $t$-values of all possible combination of paired-difference tests of the form Equation (SM6.1). The critical value $t_{\alpha=0.05,6} = 1.943$.*

| | (1,2) | (1,3) | (1,4) | (1,5) | (2,3) | (2,4) | (2,5) | (3,4) | (3,5) | (4,5) |
|---|---|---|---|---|---|---|---|---|---|---|
| $t$-value | -0.7592 | 0.6669 | 0.0879 | 0.2325 | 1.2058 | 1.124 | 1.492 | -0.2887 | -0.3973 | 0.0931 |

**SM7. Simple Linear Model.** In SM11, we report the range of each parameter for each donor and whether it was identifiable, practically unidentifiable, or structurally unidentifiable, under the assumption it takes one simultaneously bound Ab to immobilize a virion ($T = 1$) and the Ab-mucin binding rate is not affected by immobilization ($c = 1$).

**SM8. Full Model.** Let $\hat{\theta}$ denote the numeric estimate for $\vec{\theta} \in \Theta$ and $\hat{\theta}_{\text{slow}}$ denote the numeric estimate for $\vec{\theta}$ in the restricted parameter space $\Theta_{0.05,5} \cap \Theta_{\text{slow}}$. In SM12, we report

**Table SM11**

*Parameter Identifiable assuming that $T = 1$ and $c = 1$ where ID indicates a structurally identifiable parameter and PU refers to a practically unidentifiable parameters. The confidence interval are given for each parameter where $\theta_{\alpha=0.95,3} = 7.814728$.*

| | $[A]_0$ ($\mu$g/mL) | $N_*$ | $q$ | $\min(\chi^2(\vec{\theta}))$ |
|---|---|---|---|---|
| Donor F02 | ID, $[0.006, 0.04]$ | ID, $[140, 560]$ | ID, $[0.89, 0.969]$ | 5.2106 |
| Donor F05 | ID, $[0.015, 0.085]$ | ID, $[60, 260]$ | ID $[0.76, 0.908]$ | 1.6292 |
| Donor F08 | ID, $[0.020, 0.115]$ | ID, $[80, 360]$ | ID $[0.86, 0.954]$ | 0.2755 |
| Donor F13 | ID, $[0.020, 0.075]$ | ID, $[50, 100]$ | PU $[0.93, 1)$ | 15.3627 |
| Donor F15 | ID, $[0.003, 0.01]$ | ID, $[310, 780]$ | ID $[0.93, .963]$ | 4.6863 |
| Donor F17 | ID, $[0.025, 0.095]$ | ID, $[80, 210]$ | ID $[0.90, .969]$ | 3.661 |
| Donor F21 | ID, $[0.035, 0.195]$ | ID, $[70, 250]$ | ID $[0.92, .989]$ | 1.277 |

the range of each parameter for each donor within $\Theta_{0.05,5} \cap \Theta_{\text{slow}}$ and whether it was identifiable, practically unidentifiable, or structurally unidentifiable. We display the proportion immobilized predicted by our model assuming $\hat{\theta}_{\text{slow}}$ is the numeric estimate of the parameters restricted to the subspace $\Theta_{0.05,5} \cap \Theta_{\text{slow}}$ for each Donor in Figure SM13.

**Table SM12**

*The values of $T$, $c$, and $N_*$ that can permit $\vec{\theta} \in \Theta_{0.05,5} \cap \Theta_{slow}$. Lower and upper bounds of the tested parameter range are underlined. Note that $\chi^2(\alpha = 0.95, 5) = 11.0705$.*

| | $T$ | $c$ | $N_*$ | $\chi^2(\hat{\theta}))$ | $\chi^2(\hat{\theta}_{\text{slow}})$ |
|---|---|---|---|---|---|
| Donor F02 | [19,$\underline{60}$ | $[44.67, \underline{10^4}]$ | [120,620] | 0.7070 | 9.1772 |
| Donor F05 | [16,$\underline{60}$] | $[79.43, \underline{10^4}]$ | [90,370] | 0.3912 | 4.1879 |
| Donor F08 | [17,$\underline{60}$] | $[56.23, \underline{10^4}]$ | [100,540] | 0.1562 | 1.2343 |
| Donor F13 | {13} | $[2818.38, \underline{10^4}]$ | {70} | 10.0055 | 18.0788 |
| Donor F15 | [30,$\underline{60}$] | $[44.68, \underline{10^4}]$ | [250, 630] | 4.2209 | 4.2209 |
| Donor F17 | [17,34 | $[281.84, \underline{10^4}]$ | [100, 200] | 1.6432 | 3.0156 |
| Donor F21 | [19,27] | $[10^3, \underline{10^4}]$ | [120,170] | 0.5974 | 8.1838 |

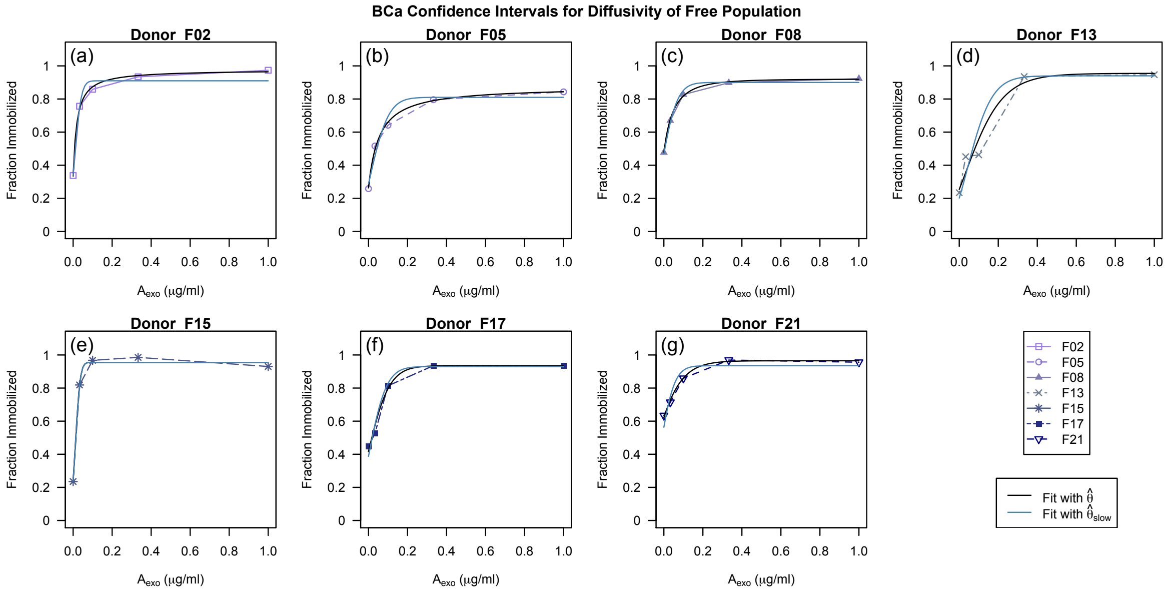


**Figure SM13.** ***(a)-(g)*** *For each donor the observed proportion immobilized (dashed lines), proportion immobilized predicted by* $\widetilde{\pi}([A];\hat{\theta})$*, (solid black line), where* $\hat{\theta}$ *is the numeric estimate for* $\vec{\theta}$ *and the proportion immobilized predicted by* $\widetilde{\pi}([A];\hat{\theta}_{slow})$ *(solid blue line)), where* $\hat{\theta}_{slow}$ *is the numeric estimate restricted to the subspace* $\Theta_{0.05,5} \cap \Theta_{slow}$*. Note in Figure (e)* $\hat{\theta}_{slow} = \hat{\theta}$*.*
